# The relaxin receptor RXFP1 signals through a mechanism of autoinhibition

**DOI:** 10.1101/2022.01.22.477343

**Authors:** Sarah C. Erlandson, Shaun Rawson, James Osei-Owusu, Kelly P. Brock, Xinyue Liu, Joao A. Paulo, Julian Mintseris, Steven P. Gygi, Debora S. Marks, Xiaojing Cong, Andrew C. Kruse

**Affiliations:** Department of Biological Chemistry and Molecular Pharmacology, Blavatnik Institute, Harvard Medical School, Boston, Massachusetts 02115, USA; Department of Systems Biology, Blavatnik Institute, Harvard Medical School, Boston, Massachusetts 02115, USA; Institut de Génomique Fonctionnelle, Université de Montpellier, CNRS, INSERM 34094 Montpellier Cedex 5, France

## Abstract

The relaxin family peptide receptor 1 (RXFP1) is the receptor for relaxin-2, an important regulator of reproductive and cardiovascular physiology. RXFP1 is a multi-domain G protein-coupled receptor (GPCR) with an ectodomain consisting of an LDLa module and leucine-rich repeats. The mechanism of RXFP1 signal transduction is clearly distinct from that of other GPCRs, but remains very poorly understood. Here, we present the cryo-electron microscopy structure of active-state human RXFP1, bound to a single-chain version of the endogenous agonist relaxin-2 and to the heterotrimeric G_s_ protein. Evolutionary coupling analysis and structure-guided functional experiments reveal that RXFP1 signals through a mechanism of autoinhibition, wherein the receptor’s extracellular loop 2 occupies the orthosteric site in the active state but is inhibited by the ectodomain in the absence of relaxin-2. Our results explain how an unusual GPCR family functions, providing a path to rational drug development targeting the relaxin receptors.

## Main

RXFP1 is a member of the leucine-rich repeat-containing GPCRs (LGR), a subset of family A GPCRs that have remained a notable exception to our understanding of GPCR signaling, despite substantial progress in studies of other GPCR families. In LGRs, the leucine-rich repeats (LRRs) function as the extracellular ligand-binding domain for three types of protein agonists: glycoprotein hormones, R-spondins, and relaxins^1^. The LGRs are involved in a variety of physiological processes across reproductive and developmental biology. RXFP1, the receptor for the relaxin-2 hormone in humans^2^, plays an important role during pregnancy. In this setting, it is responsible for physiological changes including increasing cardiac output and remodeling reproductive tissues to facilitate parturition^3–5^. RXFP1 signaling also regulates the physiology of numerous organs in both males and females, particularly the heart, lungs, liver, and kidneys. Activation of RXFP1 by relaxin-2 in these organs leads to pleiotropic cellular effects, including vasodilation, angiogenesis, anti-inflammatory responses, and extracellular matrix remodeling through collagen degradation^6,7^. Accordingly, the RXFP1 receptor has emerged as a promising therapeutic target for the treatment of cardiovascular and fibrotic diseases^8–10^.

The relaxin receptors RXFP1 and RXFP2 are unique members of the LGR family, and are classified as type C LGRs due to the presence of an additional domain called an LDLa module at the receptors’ distal N-termini, before the LRRs in sequence^11,12^. These receptors are the only two mammalian GPCRs to contain an LDLa module, and the role of this domain in relaxin receptor signaling is poorly understood. The LDLa module is dispensable for relaxin-2 binding to the LRRs but is essential for activation of RXFP1 signaling in response to relaxin-2^13^. The mechanisms that couple ligand binding in the LRRs to conformational changes within the 7TM domain required for G protein signaling remain undefined, largely due to an absence of structural data. Recent structures of the luteinizing hormone-choriogonadotropin receptor (LHCGR), one of the glycoprotein hormone receptors or type A LGRs, revealed conformational changes of the LRRs between inactive and active states. These studies proposed that large glycoprotein hormones signal through a steric “push-pull” mechanism that activates the receptor by driving changes in LRR conformation^14^. However, the small 6 kDa size of the relaxin-2 peptide precludes such a mechanism, requiring an alternative explanation.

In order to elucidate the basis for RXFP1 signal transduction, we set out to determine the active-state structure of human RXFP1 bound to an engineered relaxin-2 and the heterotrimeric G protein G_s_. We optimized receptor expression using fusions to a minimal Ga protein^15^, then formed a larger complex with the addition of G protein β_1_ and γ_2_ subunits, and determined the structure using cryo-electron microscopy. Unexpectedly, the structure revealed that RXFP1’s extracellular loop 2 (ECL2) occupies the GPCR orthosteric ligand-binding pocket in the active state. Results from structural and functional studies define a mechanism in which ECL2 conformation is regulated by the receptor’s LRRs and hinge region, a short segment between the LRRs and 7TMs. Collectively, these studies identify several conformational switches in both the receptor ectodomain and 7TMs, showing that the concerted action of multiple receptor domains controls the transduction of RXFP1 signaling by its agonist, relaxin-2.

## Results

### Cryo-EM structure of the RXFP1–G protein complex

Wild type (WT) full-length RXFP1 receptor could be expressed only at very low levels in mammalian cells. To enable structural studies, we cloned a fusion of RXFP1 to the engineered Ga protein minimal G_s_ (mini-G_s_)^15^. Truncations of RXFP1’s flexible C-terminus further increased expression levels, with the optimal expression construct having a C-terminal truncation of 20 amino acids (**Fig. S1**). The binding of the catalytically inactive mini-G_s_ protein blocks RXFP1 signaling through endogenous G_s_ proteins and likely stabilizes the receptor, leading to higher purification yields. The fusion protein of RXFP1 and mini-G_s_ was purified in complex with human G protein β_1_ and γ_2_ subunits, the camelid antibody VHH fragment nanobody 35 (Nb35)^16^, and an engineered version of the agonist relaxin-2 (SE001)^17^ to form an agonist–GPCR–G protein complex (hereafter referred to as RXFP1–G_s_).

Cryo-electron microscopy (cryo-EM) was used for structural studies of RXFP1–G_s_. Initial two-dimensional classification analysis revealed averages that showed clear density for RXFP1’s 7TM domain and the heterotrimeric G protein. In contrast, the density for RXFP1’s ectodomain was weak and poorly defined, indicating flexibility of the ectodomain with respect to the transmembrane domain. Due to the conformational heterogeneity of RXFP1–G_s_, we analyzed the cryo-EM data using two different approaches. The first utilized masking of RXFP1’s 7TM and G proteins to obtain a high-resolution cryo-EM map of these domains at 3.2 Å, allowing us to build an atomic model (**Fig. S2**). The second approach used focused classifications of RXFP1’s ectodomain to obtain a cryo-EM map of the entire complex (**Fig. S3**). As a result of the ectodomain’s flexibility, the cryo-EM map of the full-length receptor is lower resolution, with an overall resolution of 4.2 Å and local resolution of the ectodomain between 5-8 Å.

RXFP1’s 7TM domain displays characteristic hallmarks of the active state for family A GPCRs (**Fig. 1**). Most notable is the outward conformation of the intracellular end of transmembrane helix 6 (TM6), which creates the binding site for the α5 helix of Gas^18,19^. Additional active-state features include an open conformation of the “ionic lock” between Glu623^6.30^ and Lys510^3.50^ and the hydrogen bond between Tyr681^7.53^ of the conserved NPxxY motif and Tyr599^5.58^ (superscript indicates Ballesteros-Weinstein numbering system)^20^. Hydrogen bonds between Tyr681^7.53^ and Tyr599^5.58^ in active-state family A GPCRs are coordinated by a bridging water molecule, not visible at the resolution of our map^19^. In RXFP1–G_s_, the active state of the 7TMs displays a canonical interaction with heterotrimeric G_s_, similar to previously reported GPCR–G protein structures^21^.

**Fig. 1.**
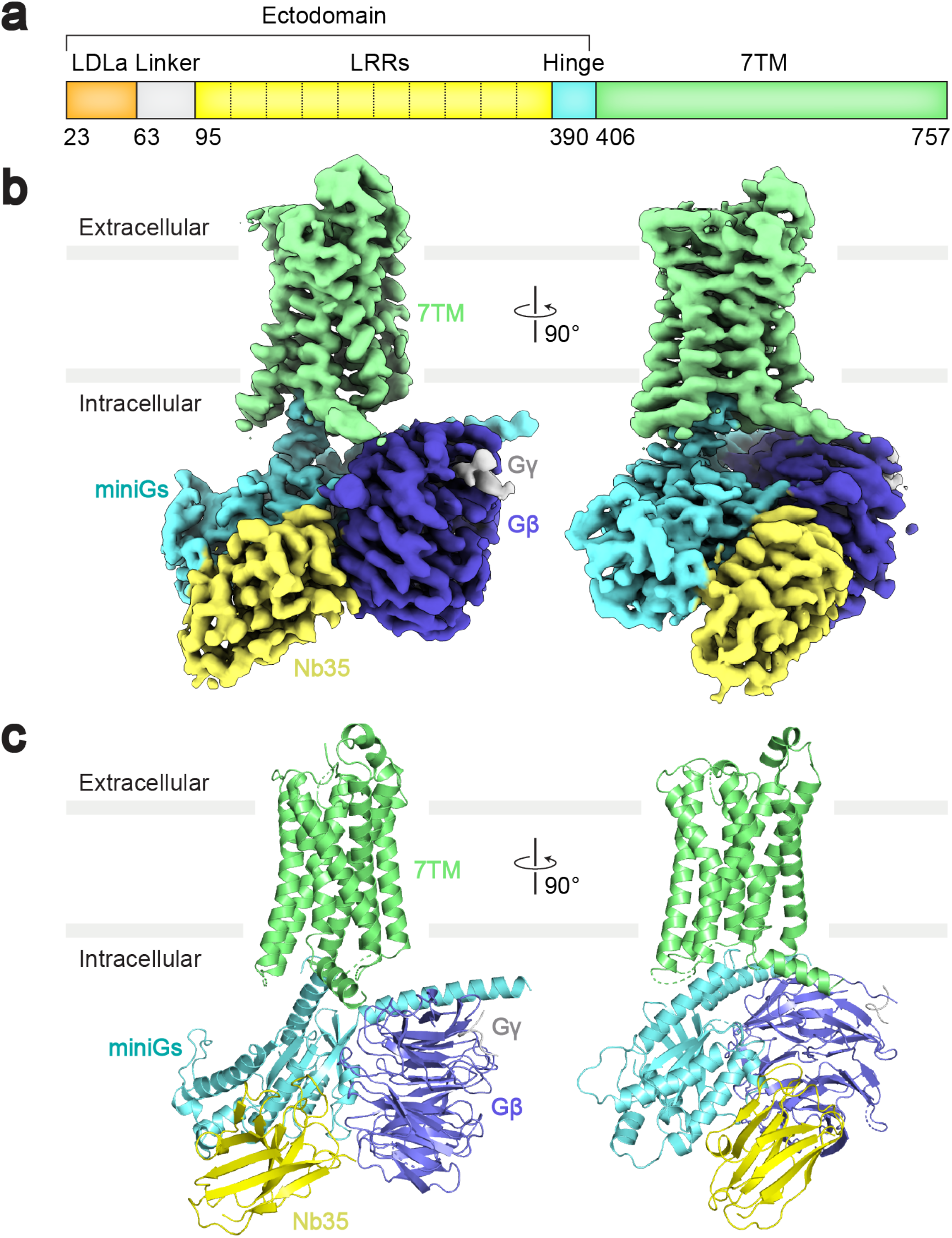
Cryo-EM map and model of the RXFP1–G_s_ complex. **a,** Diagram of the primary structure of RXFP1 domains. **b-c**, Cryo-EM map (**b**) and model (**c**) of the RXFP1–G_s_ complex 7TM domain with heterotrimeric G_s_ proteins and Nb35.

### Extracellular loop 2 is essential for signaling

The active-state structure of RXFP1 surprisingly revealed that ECL2 occupies the GPCR orthosteric ligand-binding pocket (**Fig. 2a**). The conformation of ECL2 was heterogeneous, and three-dimensional focused classifications of RXFP1’s extracellular loops were required to obtain a cryo-EM map suitable for model building. The structure of ECL2 can be described in three segments. The first segment of ECL2, from Lys550^ECL2^ to Gly558^ECL2^, interacts with TMs 3, 4, and 5, with the residues Tyr556^ECL2^ and Tyr557^ECL2^ making the most extensive contacts within the 7TMs. The second segment of ECL2, beginning at Thr559^ECL2^ until His567^ECL2^, forms a loop structure which binds into the canonical GPCR orthosteric binding site. In particular, the side chains of the residues Phe564^ECL2^ and Leu566^ECL2^ fit into a hydrophobic cavity created by TMs 2, 3, 5, 6, and 7 (**Fig. 2b**). Within this segment, Cys563^ECL2^ forms a disulfide bond with Cys485^3.25^ in TM3, a highly conserved feature among family A GPCRs which stabilizes the conformation of the loop^22^. As a result of the disulfide bond and interactions with the hydrophobic cavity, the second segment is the most well-resolved region of ECL2. In contrast, four residues in the third segment of ECL2, from Ser568^ECL2^ to Ser573^ECL2^, are not visible in the cryo-EM map, likely indicating a region of higher intrinsic flexibility.

The unusual conformation of ECL2 and deeply buried positions of Phe564^ECL2^ and Leu566^ECL2^ suggested that they may mimic the role played by exogenous agonists in other receptors. Indeed, the location of the residues Phe564^ECL2^ and Leu566^ECL2^ corresponds with the binding sites of small molecule and peptide orthosteric agonists of other family A GPCRs, such as adrenaline binding to the β_2_ adrenergic receptor^23^ and angiotensin II analogs to the angiotensin II type I receptor^24^ (**Fig. S4b,c**). Based on these observations, residues from the second segment of ECL2 were tested by mutagenesis for their contribution to signaling. ECL2 substitutions maintained above 50% of wild type RXFP1 expression, excepting the mutation of Leu566^ECL2^ to Asp, which expressed at 33% of wild-type levels (**Fig. S5a**). Mutation of Phe564^ECL2^ to Ala or Leu566^ECL2^ to Asp almost completely ablated RXFP1 signaling in response to relaxin-2, confirming the importance of these residues to receptor activation (**Fig. 2c**). Mutation of the Pro565^ECL2^ residue between Phe564^ECL2^ and Leu566^ECL2^ also decreases relaxin-2 signaling, likely by disrupting the loop structure of these ECL2 residues within the orthosteric site (**Fig. 2c**)^25^.

The discovery of the importance of ECL2 to RXFP1 signaling is reminiscent of the orphan receptor GPR52, which is directly activated by its own ECL2 as a tethered agonist^26^. While ECL2 shows no conservation of sequence or detailed structure between RXFP1 and GPR52, the binding sites for ECL2 are in analogous positions within the 7TM domain for both GPCRs (**Fig. S4a**). These structural parallels, along with RXPF1 mutagenesis in cell signaling assays, are consistent with ECL2 serving a critical role in activating RXFP1.

**Fig. 2.**
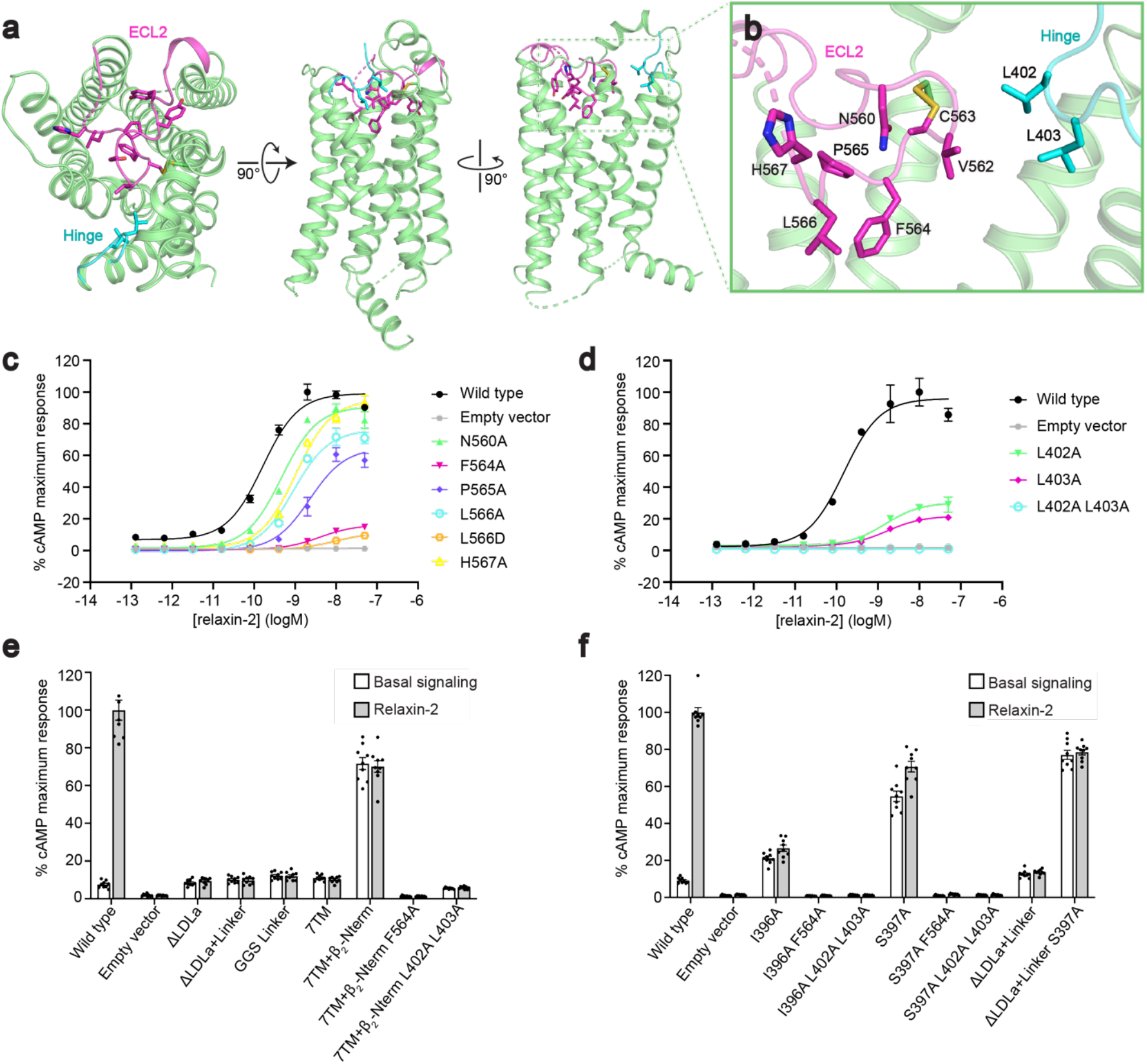
Regulation of receptor signaling by ECL2 and the ectodomain. **a,** The conformation of ECL2 and the hinge region in active-state RXFP1. **b**, Details of ECL2 in the 7TM orthosteric site and interactions between ECL2 and the hinge region. **c-d**, The effect of ECL2 (**c**) and Leu402 and Leu403 hinge region (**d**) mutations in an assay for G_s_ signaling by RXFP1. Data are mean ± s.e.m. from technical triplicates. **e-f**, Basal signaling and signaling in response to 50 nM relaxin-2 for RXFP1 ectodomain truncation constructs (**e**) and Ile396 and Ser397 hinge region mutations (**f**) in assay for G_s_ signaling. Data are mean ± s.e.m. from nine technical replicates.

### The role of the hinge region in receptor activation

Our structure of the 7TM domain also includes 6 residues from the hinge region of RXFP1’s ectodomain. The hinge region is composed of residues between the end of the LRRs and the beginning of TM1, and the six residues adjacent to TM1 are included in our refined model. We observed that the hinge region curves into the top of the 7TM domain, with residues Leu402 and Leu403 in close proximity to ECL2 in RXFP1’s orthosteric site. Additionally, residues from both ECL2 and the hinge region pack against the top of TM7 (**Fig. 2b**).

Functional analyses of type A LGRs, the glycoprotein hormone receptors (GPHRs), have previously established that the hinge region of those LGRs is critical for receptor signaling^27,28^. The unstructured loop in the hinge region of GPHRs is approximately 60-125 residues long and plays a role in binding the receptors’ glycoprotein hormone agonists^29^. For GPHRs, a 10-residue section of their hinge region near the 7TM domain, P10, is also critical for receptor signaling^27,28^ and has been recently shown to adopt different conformations in inactive and active receptor states of LHCGR^14^. In contrast, the hinge loop of type C LGRs, such as RXFP1, is predicted to be about 15 residues long, has no established function, and shows no sequence conservation in comparison to type A LGRs^1^. Despite the clear differences between GPHRs and RXFP1, the interactions between the hinge region and orthosteric site in RXFP1’s active-state structure unexpectedly suggested that these residues may also play a role in signaling.

To investigate the role of the hinge region residues Leu402 and Leu403, we cloned single mutations to Ala and constructed a double mutant with Ala substitutions for both residues. These constructs expressed at low levels, so wild type receptor expression was reduced in order to compare receptor signaling (**Fig. S5b**). While the single Ala mutants each decreased the efficacy of relaxin-2, the double Ala mutant completely ablated RXFP1 signaling, despite maintaining the ability to bind relaxin-2 and expression levels at roughly 50% of wild type (**Fig. 2d, S5f**). These results indicated that Leu402 and Leu403 of the hinge region are essential for RXFP1 activation, likely functioning to stabilize ECL2 into its active-state conformation in the orthosteric site.

### Autoinhibition of RXFP1 signaling

While structures of RXFP1 and GPR52 revealed that ECL2 is critical for the activation of both receptors, other aspects of their signaling suggest differing mechanisms. GPR52 is a self-activated orphan GPCR with very high basal activity, signaling at 90% of its E_max_ without any agonist bound^26,30^. The mechanism governing the intrinsic activity of GPR52 is clear, as the ECL2 tethered agonist is a component of the 7TM structure itself. In contrast, RXFP1 does not have high basal activity, but signals in response to the binding of relaxin-2 to its LRRs. These differences suggest that while both GPCR structures show ECL2 binding in the orthosteric site, RXFP1 likely uses additional mechanisms to prevent continuous self-activation of the 7TM domain. The major structural difference between these two receptors is that GPR52 is a conventional family A GPCR with an unstructured N-terminus, while RXFP1 is a type C LGR, containing a structured ectodomain of LRRs and an LDLa module. For this reason, we set out to investigate the role of RXFP1’s ectodomain in modulating the activity of ECL2.

To address this question, we first cloned constructs of RXFP1 with deletions of the receptor’s ectodomain. Basal signaling of RXFP1 was not increased by deletion of the LDLa module, deletion of the LDLa module and the 32-residue linker that connects it to the LRRs, or replacement of the linker with a 32 residue Gly-Gly-Ser linker (**Fig. 2e**). These results indicated that the LDLa module and linker of RXFP1’s ectodomain do not play an inhibitory role in signaling. In contrast, these constructs are unable to signal in response to relaxin-2, consistent with previous studies showing that the LDLa module and linker region are essential for receptor activation and that the linker is involved in relaxin-2 binding^13,31^ (**Fig. 2e, S5e**). We next focused on testing the role of RXFP1’s LRRs in the receptor signaling mechanism. To remove the ectodomain, including the LRRs, constructs were designed to express RXFP1’s 7TM domain alone, which also included several residues from the hinge region immediately preceding TM1 (**Table S8**). However, these constructs showed very low cell-surface expression, at 12% of wild type. Although the 7TM domain expressed very poorly, it showed similar basal signaling to wild type RXFP1 in a G_s_ signaling assay, at 11% of the wild type relaxin-2 E_max_. To increase expression of the 7TMs, we cloned a fusion of RXFP1’s 7TM domain to the unstructured N-terminus of the high-expressing β_2_ adrenergic receptor. This fusion rescued 7TM domain expression to essentially wild type levels (**Fig. S5c**). When tested in a G_s_ signaling assay, the fusion showed a high level of basal activity. In the absence of ligand, the 7TM domain signaled at 70% of the maximum level of agonist-induced signaling for wild-type RXFP1 (**Fig. 2e**). As expected, RXFP1’s 7TM domain alone does not show any change in activity in response to relaxin-2, since the relaxin-2 binding sites in the ectodomain are deleted in this construct^32^ (**Fig. 2e, S5f**).

To establish whether the high basal activity of RXFP1’s 7TM alone is due to ECL2, we introduced the Phe564^ECL2^ to Ala mutation that greatly reduced full-length RXFP1 signaling in response to relaxin-2. The Phe564^ECL2^ to Ala mutation was able to completely ablate the 7TM domain’s high basal activity, confirming that ECL2 constitutively activates the 7TMs in the absence of RXFP1’s LRR domain. Likewise, Ala mutations of the hinge region residues Leu402 and Leu403 (which are included in the 7TM domain fusion construct) were also able to reduce the high basal signaling. These data suggested a model of signaling in which the LRRs play an autoinhibitory role in regulating the active state of ECL2 and the hinge region. As a result, deletions of the LRRs allow constitutive activation the receptor, leading to high basal signaling.

### Inhibitory interactions revealed by evolutionary coupling analysis

An additional insight into the regulation of ECL2 by the ectodomain arose from evolutionary coupling (EC) analysis of RXFP1^33,34^. The strongest ECs, or evolutionary coupled residues, are derived from applying a global probability model to multiple sequence alignments and typically indicate residue pairs that are in contact in 3D^35^, including residues involved in conformational changes^36,37^. Residues of ECL2 had strong ECs pairing them with residues within the 7TM helices, supporting our active-state structure. However, ECL2 also showed ECs with residues from RXFP1’s hinge region not present in our model (**Fig. S6**). In total, two residues from the hinge region, Ile396 and Ser397, had ECs with three residues of ECL2, including the critical Phe564^ECL2^. ECL2 and the hinge region are likely involved in other interactions than those observed in our structure of the active state. If ECL2 contacts Ile396 and Ser397 in an inactive-state conformation, we predicted that those interactions may contribute to the inhibition of ECL2 in the absence of relaxin-2 binding to the LRRs.

Ile396 and Ser397 were each mutated to Ala to test the effects on the activation state of ECL2. Despite having low expression levels, the Ile396 and Ser397 mutations each showed a significant increase in basal signaling, at 21% and 55%, respectively, of wild type RXFP1’s E_max_ (**Fig. 2f**, **S5d**). The Ser397 to Ala mutant similarly increased the basal signaling of RXFP1 with a deletion of the LDLa module and linker, confirming that those domains are not involved in the mechanism of signaling inhibition. Interestingly, the hinge mutants showed a reduced signaling response to relaxin-2 binding, suggesting that these residues involved in a potential inactive-state ECL2 interface are also important for allosteric communication between the ectodomain and 7TM domain (**Fig. 2f, S5f**). Addition of the Phe564^ECL2^ or Leu402 and Leu403 substitutions to the Ile396 and Ser397 mutants was able to abolish the increase in basal signaling, confirming that RXFP1 activation is dependent on these residues (**Fig. 2f**).

### Mechanism of RXFP1 7TM autoactivation

We performed enhanced-sampling molecular dynamics (MD) simulations (see Methods for details) to study the role of ECL2 and the hinge region in the basal activity of RXFP1’s 7TM domain. An inactive-state model of the 7TM domain was obtained by MD simulations of deactivation, starting from the cryo-EM structure truncated before the hinge region (residues 395–699). A sodium ion was placed in the sodium-binding site to favor the sampling of inactive states during the simulations. We obtained an inactive state that was strikingly similar to the inactive-state AlphaFold2 model^38^, in which the intracellular end of TM6 bent toward TM3 to form the classic Lys510^3.50^–Glu623^6.30^ ionic lock (**Fig. S7a-c**). The second segment of ECL2 maintained the same conformation throughout the simulations, whereas Ser568^ECL2^ to Ser573^ECL2^ in the third segment were highly mobile, consistent with absence of clear density in the cryo-EM map. Starting from the inactive state, we first simulated autoactivation of the RXFP1 7TM by removing the sodium ion and protonating the sodium anchor, Asp451^2.50^. As negative controls, we performed the same simulations for two mutants with low basal activity, F564^ECL2^A and L566^ECL2^D. The WT RXFP1 7TM exhibited autoactivation with outward movements of TM6 on the intracellular side, destabilization of the ionic lock, and frequent side-chain flips of the toggle switch W641^6.48^ (**Fig. 3a-c**). In contrast, the two mutants remained in inactive conformations. The WT 7TM exhibited distinct shapes of the orthosteric pocket compared to the two mutants, owing to ECL2–TM7 interactions. Namely, Phe564^ECL2^ in the WT 7TM domain “pushed” TM7 toward TM2, which likely altered W641^6.48^ conformations and triggered autoactivation (**Fig. 3a-c**). The F564^ECL2^A mutation directly eliminates this steric effect, whereas substituting Leu566^ECL2^ with Asp reorients the charged side chain away from the pocket, leaving space for F564^ECL2^ and diminishing its impact on TM7 conformation (**Fig. 3a,b**).

The truncated 7TM model is unsuited for studying the hinge region, which partially unfolds during the simulations. Therefore, we used a truncated ^half^LRRs-7TM form (residues 242–699) to investigate the role of the hinge. The ^half^LRRs-7TM model was truncated before the Cys243-Cys279 disulfide bond in the middle of the LRRs to prevent unfolding, while reducing the system size to enable sufficient MD sampling. MD simulations were performed for the constitutively active mutant S397A, in comparison with the WT and the triple mutant S397A/L402A/L403A, starting from the inactive state. We found that the S397A mutation disrupts H-bonds in the hinge and increases the mobility of the LRRs, which diminishes the autoinhibition of ECL2 and promotes 7TM activation (**Fig. 3d and Table S6**). L402A/L403A attenuates the effect of S397A by stabilizing the receptor in a different conformation (**Fig. 3d**).

**Fig. 3.**
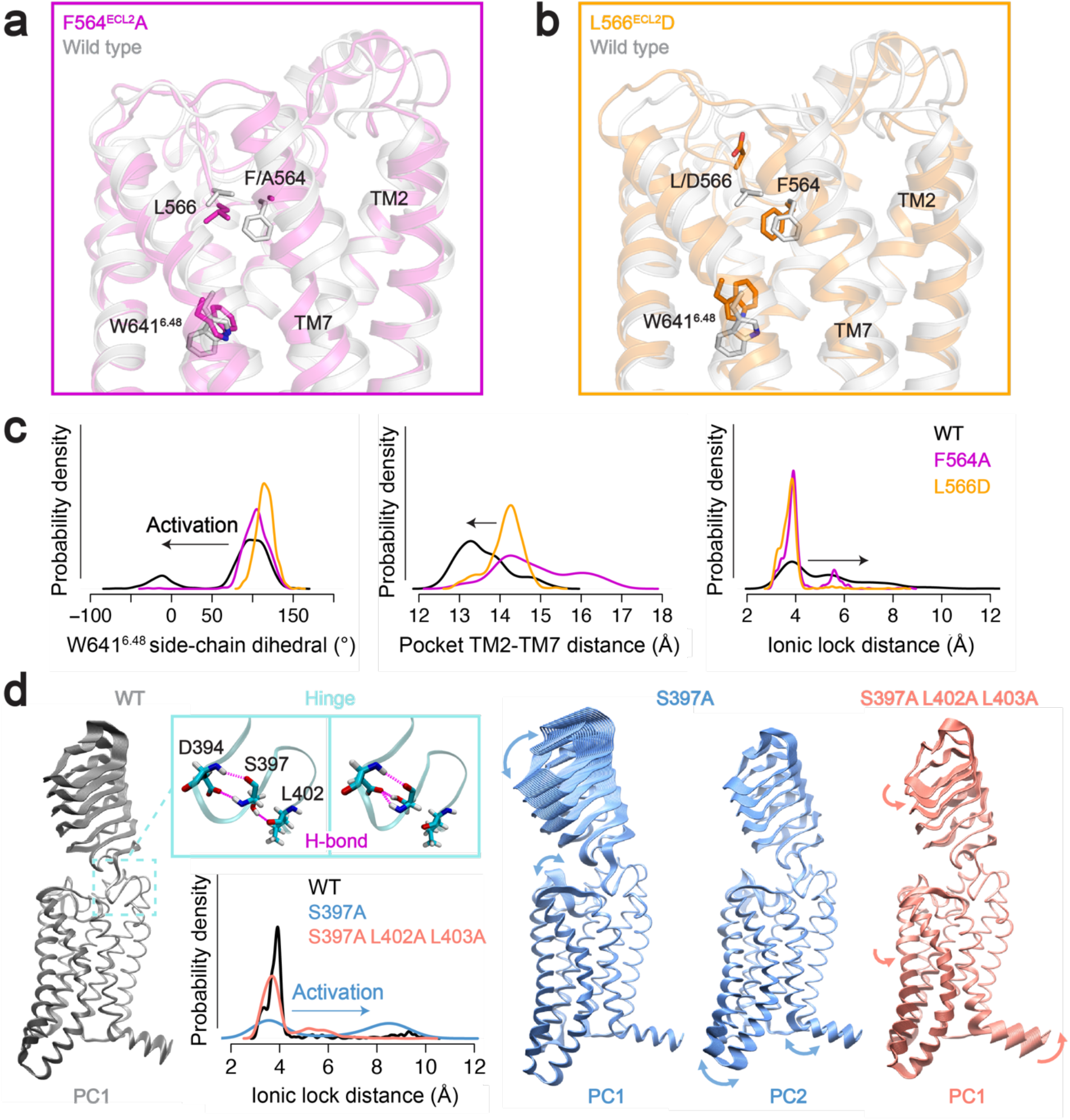
Molecular dynamics of RXFP1 starting from the inactive-state AlphaFold2 model. **a-b,** The truncated RXFP1 7TM domain alone shows autoactivation. Autoactivation in these simulations is impaired by the addition of the F564A (**a**) or the L566D mutations (**b**). **c**, Histograms describing activation-related conformational differences between WT, F564A, and L566D RXFP1 7TM models, including the distance between TM2 and TM7 in the orthosteric site, side-chain flips of the toggle switch residue W641^6.48^, and the ionic lock distance. **d,** Molecular dynamics of truncated RXFP1 ^half^LRRs-7TM. Projection of the trajectories on the first and second principal components (PC) illustrates the mechanism of S397A-induced basal activity. The S397A mutation disrupts the H-bonds with L402 (backbone) and D394 (side chain) present in the WT (Table S6). The hinge and LRRs become more mobile in the S397A mutant, which triggers activation through ECL2. Addition of the L402A/L403A mutations reduces the steric hindrance of the hinge and leads to an overall twist of the receptor, which attenuates the activation effect of S397A.

### Relaxin-2 binding to the leucine-rich repeats

The cryo-EM map of the full-length RXFP1–G_s_ complex is limited by lower resolution due to the dynamic nature of the receptor’s ectodomain. Continuous heterogeneity present in the final particle stack for RXFP1–G_s_ was visualized through three-dimensional variability analysis in cryoSPARC^39^. The resulting movies showed that the LRRs are very flexible in the relaxin-2–bound state, moving at the hinge region between the LRRs and 7TM domain (**Movie S1**). Despite these limitations, the cryo-EM map offered several new insights into the overall ectodomain architecture and relaxin-2 binding.

In the refined cryo-EM map of active-state RXFP1, the LRRs are positioned above ECL1, likely giving this region a role in stabilizing ectodomain conformations. Mutation of the ECL1 residue W479^ECL1^ has been previously shown to reduce both relaxin-2 binding and signaling^25^. In the active-state structure, W479 projects from ECL1 to interact extensively with residues in the 7TM domain. Most or all of these interactions would be abrogated by the W479A substitution, accounting for lack of RXFP1 function due to structural changes that would destabilize the receptor.

The LRRs are extended away from the transmembrane domain in the active state, at an angle of 40° from the membrane plane (**Fig. 4a**). The orientation of the ectodomain also rotates the concave ligand-binding side of the LRRs away from the extracellular side of the 7TM domain (**Fig. 4b**). Each of these features physically separates the relaxin-2 binding site on the LRRs from the 7TMs. While a secondary binding site between relaxin-2 and the extracellular loops (ECLs) has been proposed^25,40^, the active state that we captured through cryo-EM does not show any direct interaction between relaxin-2 and the ECLs.

The low-resolution map shows relaxin-2 bound to the concave side of the LRRs. To aid with modeling the relaxin-2**–** LRR interaction, we used crosslinking mass spectrometry (CLMS) with the RXFP1**–** G_s_ complex. Using the Extended-EDC approach with an EDDA crosslinker^41^, Glu14^B-chain^ of relaxin-2 crosslinked with three residues on the LRRs, Glu206, Glu299, and Glu351 (**Fig. 4c,d**). Several residues involved in relaxin-2 binding have been previously characterized through mutational analysis in radioligand binding assays. In those experiments, a series of conserved residues on the B-chain of relaxin-2, Arg13, Arg17, and Ile/Val20, a motif known as the relaxin binding cassette, were proposed to be part of the interface^42,43^. Multiple residues on the concave side of the LRRs were also found to be important for relaxin-2 binding^32^. Interestingly, the Glu14^B-chain^ residue of relaxin-2 crosslinked to the LRRs is adjacent to the Arg13^B-chain^ of the relaxin binding cassette. Additionally, Glu299 on the LRRs had been previously identified as a residue involved in relaxin binding by mutational studies, highlighting the close agreement between our CLMS data and prior functional analyses.

The residues from our CLMS experiment were used in combination with the mutational data as restraints for docking the relaxin-2–LRR interaction in HADDOCK^44^ (**Fig. 4c,d**). The highest scoring HADDOCK models agreed well with our low resolution cryo-EM map and showed the B-chain of relaxin-2 bound to the concave side of the LRRs, while the A-chain made limited contacts (**Fig. 4e**). Additional density in the cryo-EM map is present at multiple sites of potential N-linked glycosylation and next to the A-chain **(Fig. S8e,f**). Density near the A-chain of relaxin-2 may belong to the ectodomain’s linker region, which the A-chain has been proposed to bind^31^. However, the low resolution and absence of crosslinks for these domains prevented further characterization of A-chain interactions.

In the docked model of relaxin-2 bound to the LRRs, Glu14^B-chain^ falls within the E-EDC crosslink distance of 14 Å from Glu206, Glu299, and Glu351 and is not directly involved in the relaxin-2 binding interface, in agreement with previous mutational analysis^43^. The model also predicted that one of the relaxin binding cassette residues, Arg17^B-chain^, interacts with Glu206, a residue on the LRRs identified by CLMS that was not previously known to be involved in the binding site. To verify this interaction, we mutated Glu206 to Ala and tested the effect of the mutation on binding of an Fc-tagged relaxin-2 protein (SE301)^17^. The Glu206 to Ala mutant expressed at equivalent levels to wild type receptor, but the single mutant reduced relaxin-2 binding to 42% of wild type levels (**Fig. 4f,g**), validating the proposed interaction.

**Fig. 4.**
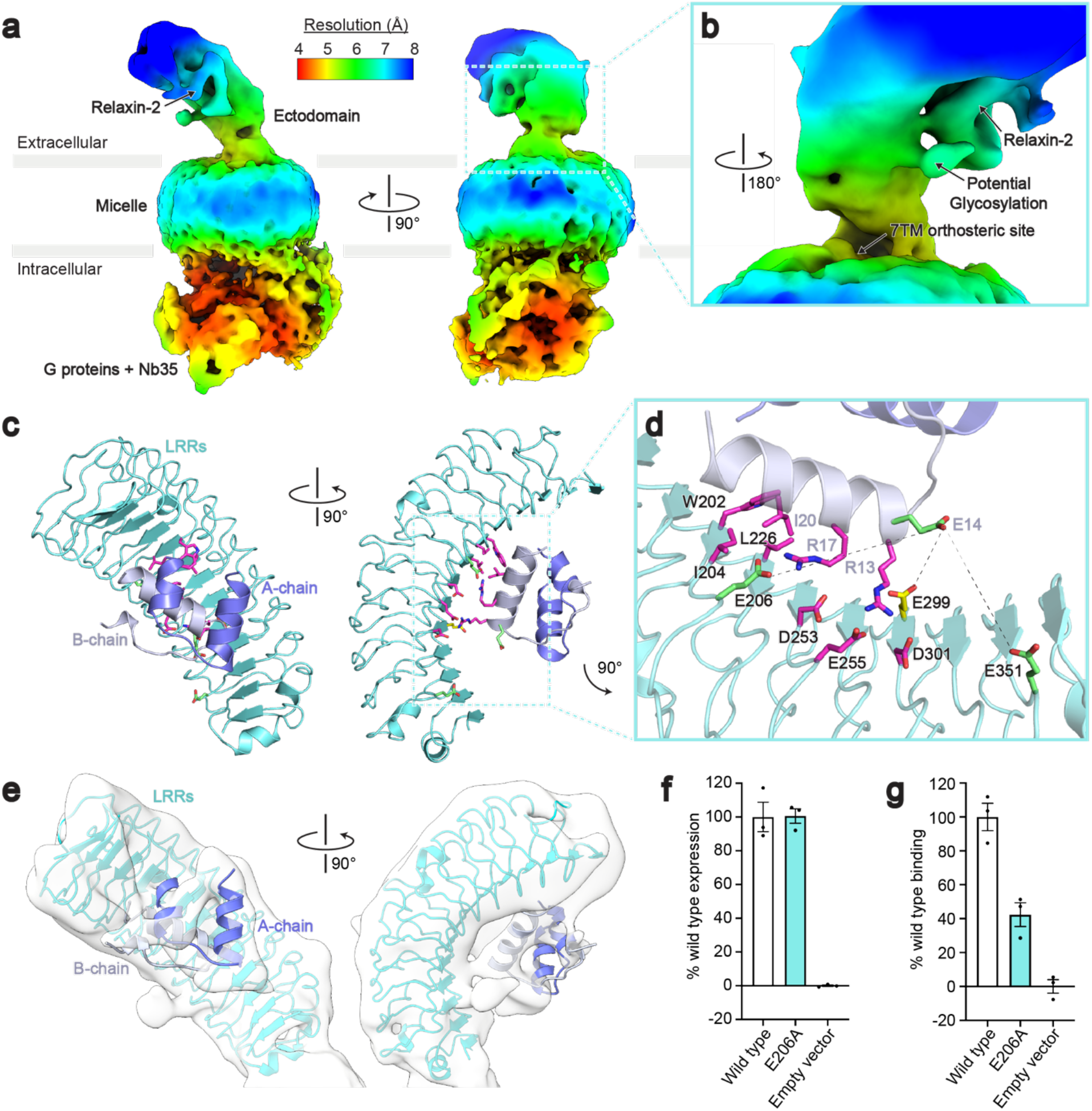
Cryo-EM and crosslinking mass spectrometry reveal interactions between relaxin-2 and the leucine-rich repeats. **a**, Local resolution cryo-EM map of the full-length RXFP1–G_s_ complex. **b**, The relaxin-2 binding site is above and rotated away from the 7TM orthosteric site. **c**, Model of the relaxin-2–LRR interaction from HADDOCK. **d**, Details of the relaxin-2–LRR interface with residues identified in published binding studies in magenta, residues from CLMS in green, and Glu299 from both CLMS and published binding studies in yellow. Crosslink distances: Glu14^B-chain^–Glu206 = 14.6 Å, Glu14^B-chain^–Glu299 = 10 Å, Glu14^B-chain^–Glu351 = 11.4 Å **e**, The relaxin-2–LRR model fit into the low resolution cryo-EM map. **f-g**, Receptor expression (**f**) and Fc-tagged relaxin-2 binding data (**g**) for the Glu206 to Ala mutation. Data are mean ± s.e.m. from technical triplicates.

## Discussion

Our active-state structure of RXFP1–G_s_ revealed unexpected features involved in the activation of RXFP1. Most strikingly, RXFP1’s ECL2 was observed to occupy the GPCR orthosteric ligand binding site, with the key residues Phe564^ECL2^ and Leu566^ECL2^ playing an essential role in receptor activation. Residues Leu402 and Leu403 of the hinge region were also required for RXFP1 signaling. These two signaling motifs were shown to be autoinhibited in the absence of relaxin-2 binding by both the LRRs and two residues of the receptor hinge region, Ile396 and Ser397, with the minimal inhibitory ectodomain construct requiring both of these features. We also defined the binding site of relaxin-2 on the LRRs, showing that it is physically separated from the 7TM domain in the active state.

Our observations indicate that RXFP1 cannot be controlled by the steric occlusion mechanism proposed for other GPHRs^14,45^. In fact, owing to its small size (6 kDa), relaxin-2 binding to the RXFP1 ectodomain is fully compatible with a membrane-proximal inactive conformation like that observed for the inactive-state of LHCGR (**Fig. 5a**). As a result, purely steric effects cannot drive receptor activation in RXFP1, necessitating an alternative mechanism. Moreover, RXFP1 and LHCGR have opposing LRR orientations in their active-state structures, highlighting the divergence of mechanisms between GPHRs and RXFP1 (**Fig. S9**). A possible mechanism for RXFP1 activation is suggested by the fact that, unlike other GPHRs, RXFP1 contains an LDLa domain which is strictly required for signaling (**Fig. 2e**)^13^. NMR studies with soluble constructs of RXFP1’s ECLs concluded that the LDLa module and residues from the adjacent linker region may interact with ECL2^25,31,46^, consistent with our observation that ECL2 serves as a key activation switch. The LDLa module was not resolved in our maps, suggesting that it is mobile in the active state of the receptor (**Fig. 5b**).

Notably, the structure of LHCGR in complex with G proteins^14^ shows a similar ECL2 conformation to that observed for the relaxin receptor, with a Phe515/Met517 pair positioned similarly to Phe564^ECL2^ and Leu566^ECL2^ in RXFP1. In fact, all LGRs share a CΦPΦ sequence motif in ECL2 (where “Φ” denotes a hydrophobic amino acid), and AlphaFold2 models of all LGRs show similar ECL2 conformations to those of LHCGR and RXFP1. This suggests that ECL2-triggered activation may be a general feature of the LGR family as a whole, inducing receptor activation in response to conformational changes of the LRRs and hinge region by either steric “push” of the ligand or, in the case of RXFP1, indirect rearrangement of the LDLa module.

The unusual features of RXFP1 signaling by relaxin-2 raises the question of whether other ligands, such as the small molecule agonist ML290^47^, activate the receptor through similar mechanisms. To answer this question, we assayed ML290’s signaling activity at wild type RXFP1 versus mutants of ECL2 and the hinge region, finding a similar dependence on these residues to relaxin-2 (**Fig. S10**). Our results indicate that small molecules are likely able to exploit the ECL2/hinge region conformational switch, although more work will be required to understand ML290’s mechanism in detail. Additional small molecule or biologic agonists could be created to exploit allostery in the receptor and either mimic ECL2-induced activation or relieve inhibitory interactions between the ectodomain and ECL2. Such molecules could be useful therapeutics for the treatment of numerous cardiovascular and fibrotic diseases, and similar approaches may be applicable to other members of the LGR family.

**Fig. 5.**
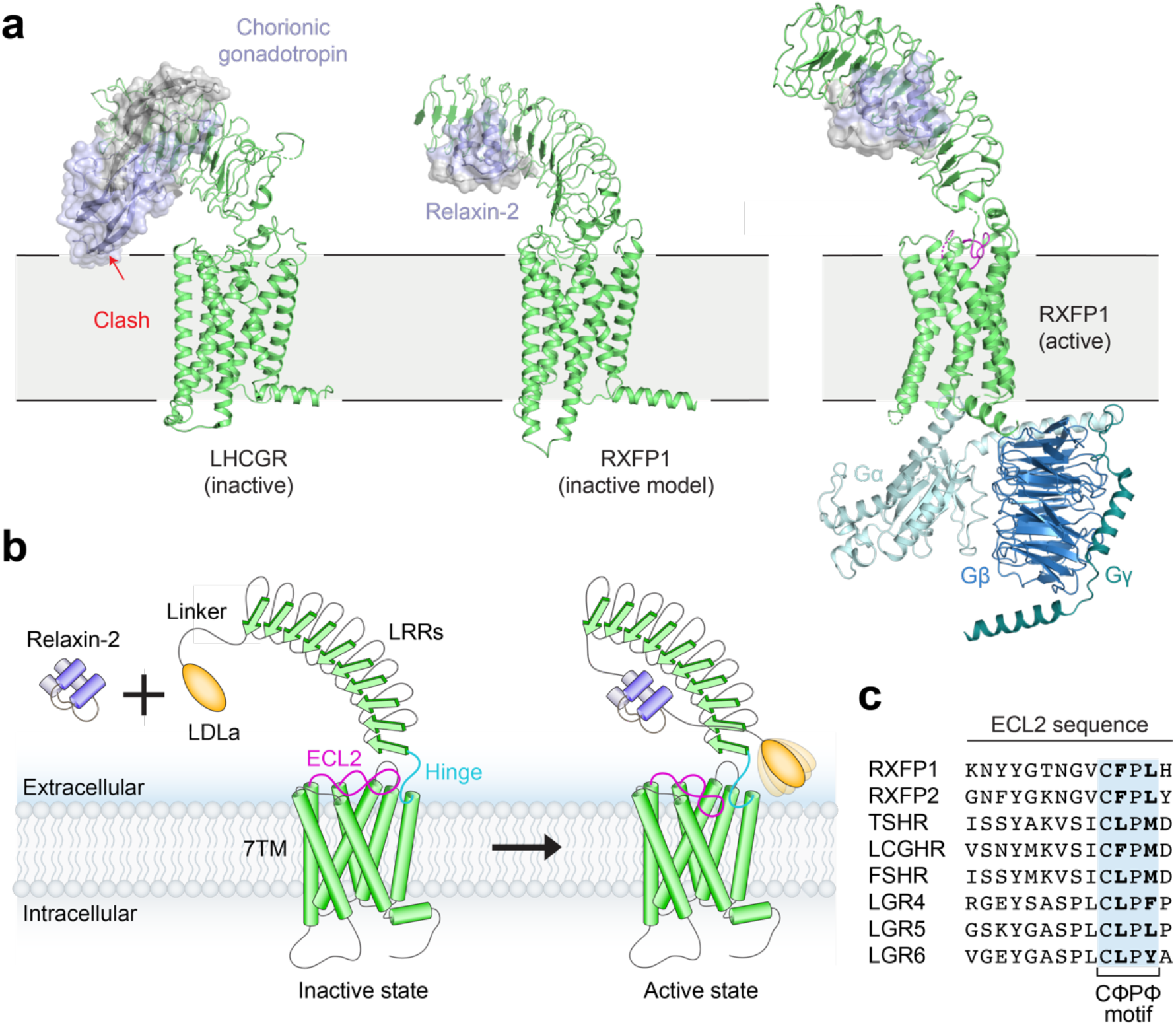
Model of RXFP1 activation by relaxin-2. **a,** Model of LHCGR in the inactive conformation (PDB ID: 7FIJ) bound to the hormone chorionic gonadotropin, showing steric clash with the membrane that induces activation^14^. In contrast, a similar potential conformation of RXFP1 would easily accommodate bound relaxin-2 (generated from alignments of inactive-state AlphaFold2 models of RXFP1’s LRRs and 7TM to PDB 7FIJ). A hybrid model of active-state RXFP1 is also shown for comparison, based on our 7TM domain structure, cryo-EM maps, and AlphaFold2 model of the LRRs with docked relaxin-2 hormone. The hybrid model uses Gγ_2_ from PDB 3SN6^16^. **b,** The inactive state of RXFP1 is characterized by inhibitory interactions between ECL2, the hinge region, and the LRRs that prevent receptor activation. Relaxin-2 binds to the concave side of the LRRs, away from the 7TM domain. Relaxin-2 binding leads to reorganization of the LDLa/hinge/ECL2 interface, allowing residues in both the hinge region and ECL2 to activate the receptor. **c,** Sequence alignment of human LGRs showing the conserved CΦPΦ motif in ECL2.

## Acknowledgements

We thank Marie Bao for critical reading of the manuscript, and the SBGrid Consortium for computational support of structural biology software. Cryo-EM data was collected at the Harvard Center for Cryo-Electron Microscopy at Harvard Medical School, and we thank them for their support and advice during data collection. This work was funded by an NIH Ruth L. Kirschstein Predoctoral fellowship (F31 GM128233) to S.C.E and a Blavatnik Biomedical Accelerator grant from Harvard Medical School to A.C.K. Molecular dynamics simulations were performed using HPC resources from GENCI-TGCC France (grants 2021-2022 Spe00015 and A0100712461).

## Data availability

The cryo-EM model and maps for RXFP1–G_s_–TM are deposited under accession codes 7TMW (PDB) and EMDB-26003, respectively. The cryo-EM map for RXFP1–G_s_–F_L_ is deposited under the accession code EMDB-26004. The rigid-body docking model of the RXFP1 LRRs bound to relaxin-2 is available on the website for the Kruse lab at Harvard Medical School.

## Author contributions

The molecular cloning, protein expression and purification, and cryo-EM grid preparation were performed by S.C.E. with supervision from A.C.K. Flow cytometry and cell signaling assays were performed by S.C.E and J.O.-O. with supervision by A.C.K. The cryo-EM data were processed by S.C.E. and S.R. Model building and refinement were performed by S.C.E. with supervision from A.C.K. The evolutionary coupling analysis was performed by K.P.B. with supervision from D.S.M. The crosslinking mass spectrometry was performed by X.L., J.A.P., and J.M. with supervision from S.P.G. The molecular dynamics simulations were performed by X.C. The manuscript was written by S.C.E., X.C., and A.C.K. with input from all authors.

## Methods

### Cloning of RXFP1 constructs

Residues 23-757 of human RXFP1 were cloned with an N-terminal hemagglutinin signal sequence, FLAG tag, and 3C protease site into the pcDNA-Zeo-tetO vector^48^. Fusions to the miniG_s_-399 protein with C-terminal truncations to RXFP1 were constructed using PCR followed by NEBuilder HiFi DNA Assembly (New England Biolabs). Truncations removed 10, 15, 20, 25, 30, or 35 residues from the receptor C-terminus. For signaling assays, human RXFP1 residues 23-757 with an N-terminal hemagglutinin signal sequence and FLAG tag were cloned into pcDNA-Zeo-tetO. Mutations to ECL2, hinge region, or LRR residues were introduced using Quikchange Lightning PCR (Agilent). Ectodomain truncations were constructed using PCR and NEBuilder HiFi DNA Assembly. Residue numbering is based on the canonical RXFP1 sequence beginning at the initiating Met residue (UniProt ID Q9HBX9).

### Cell surface expression tests

RXFP1 signaling assay constructs were tested for cell surface expression using flow cytometry. HEK293T cells (ATCC) were maintained in Dulbecco’s Modified Eagle Medium (DMEM) (Corning) with 10% (v/v) fetal bovine serum (FBS) (Sigma-Aldrich). Cells were plated at 100,000 cells/well into 12-well plates (Thermo Fisher Scientific). The following day, cells were transfected with 220 ng/well (unless otherwise stated) of human RXFP1 or empty vector DNA using FuGENE, according to the manufacturer’s instructions (Promega). For **Fig. S5b**, 4.4 ng of wild type RXFP1 DNA was used per well, plus 215.6 ng of empty vector DNA, to lower wild type receptor expression levels. For **Fig. S5c**, 110 ng of 7TM+β_2_-Nterm DNA was used per well, plus 110 ng of empty vector DNA, in order to lower the expression of this construct to be equivalent to wild type levels. After twenty-four hours, the media was aspirated, and cells were detached by pipetting in phosphate-buffered saline (PBS) with 1% (v/v) FBS and 2 mM calcium chloride (Buffer A). Cells were distributed at 100,000 cells/well in 200 μL Buffer A into a V-bottom 96-well plate (Corning). Cells were washed once and blocked in Buffer A by incubation for 30 minutes at 4°C. M1 anti-FLAG antibody (In house) labeled with Alexa Fluor 647 (Thermo Fisher Scientific) was incubated with the cells at 2.5 μg/mL for 1 hour at 4°C. Cells were washed twice in Buffer A and resuspended in 100 μL Buffer A. Fluorescence intensity was quantified using a CytoFLEX flow cytometer (Beckman Coulter). Around 2,000 events per sample were collected and analyzed using FlowJo (**Fig. S11**). Data was normalized using the wild type and empty vector mean fluorescence intensities as 100% and 0%, respectively, and plotted using GraphPad Prism.

### cAMP signaling assay

The GloSensor assay from Promega, a live-cell signaling assay that detects cellular cAMP levels, was used to measure activation of G_s_ signaling through RXFP1. White, clear-bottom 96-well plates (Thermo Fisher Scientific) were coated with 30 μL of 10 μg/mL poly-D-lysine (Sigma-Aldrich) and washed once with PBS. HEK293T cells were then plated at 2×10^4^ cells/well in DMEM with 10% (v/v) FBS. The following day, cells were transfected with 20 ng of GloSensor reporter plasmid and 20 ng (unless otherwise stated) of human RXFP1 or empty vector DNA per well using FuGENE, according to the manufacturer’s instructions. For **Fig. 2d**, 0.4 ng of wild type RXFP1 DNA was used per well, plus 19.6 ng of empty vector DNA, according to ratios determined during cell surface expression tests. For **Fig. 2e**, 10 ng of 7TM+β_2_-Nterm DNA was used per well, plus 10 ng of empty vector DNA, according to ratios determined during cell surface expression tests. Twenty-four hours later, the media was replaced with 40 μL of CO_2_-independent media (Thermo Fisher Scientific) with 10% (v/v) FBS and 2 mg/mL D-luciferin (Goldbio) and incubated for 2 hours at room temperature (RT) in the dark. Measurements of luminescence with 1 second integration times were taken before ligand addition using a SpectraMax M5 microplate reader. For signaling curves, dilution series of native relaxin-2 (R&D Systems), were added to the cells and luminescence measurements were taken 30 minutes after relaxin-2 addition. For measurements of RXFP1 basal signaling versus agonist-induced signaling, vehicle control (PBS + 0.1% bovine serum albumin), relaxin-2 at 50 nM final concentration, or ML-290 at 490 nM final concentration were added to the cells and the luminescence measurement was taken after 30 minutes. The maximum signaling response of relaxin-2 or ML-290 at wild type RXFP1 for each experiment was normalized to 100%, and the percentages were plotted using GraphPad Prism.

### Expression of RXFP1-miniG_s_

RXFP1-miniG_s_ constructs for protein purification were expressed in inducible Expi293F tetR cells^48^ (Thermo Fischer Scientific). Stable cells lines were generated for RXFP1-miniG_s_399-20res by plating Expi293F tetR cells adherently in DMEM with 10% (v/v) FBS. Cells were transfected with linearized RXFP1-miniG_s_399-20res pcDNA-Zeo-tetO DNA using Lipofectamine, according to the manufacturer’s protocols (Thermo Fischer Scientific). Stable integrations were selected using 200 μg/mL Zeocin. After selection, polyclonal adherent RXFP1-miniG_s_399-20res cells were readapted to suspension culture in antibiotic-free Expi293 media (Thermo Fischer Scientific), then maintained in Expi293 media with 10 μg/mL Zeocin. Stable RXFP1-miniG_s_399-20res Expi293F tetR cells were expanded for protein expression in antibiotic-free Expi293 media and induced with 4 μg/mL doxycycline, 0.4% glucose, and 5 mM sodium butyrate. After 48 hours of induction, cells were harvested by spinning at 4,000 xg for 30 minutes at 4°C. The pellet was flash frozen in liquid nitrogen and stored at −80°C.

### Purification of relaxin proteins

Single-polypeptide versions of relaxin-2 were used for RXFP1 complex purifications and flow cytometry binding assays. These single-polypeptide relaxin-2 constructs utilize linkers to connect relaxin-2’s B-chain and A-chain. Sequences for these proteins can be found in **Table S8** and their design and characterization are described in detail elsewhere^17^.

Single-chain relaxin proteins were purified as previously described^17^. Briefly, His-tagged single-chain relaxin-2 (SE001) was expressed as a secreted protein from inducible Expi293F tetR cells. SE001 was purified from cell supernatants after 5 days of induction using Nickel Excel resin (GE Healthcare), followed by size exclusion chromatography with a Sephadex S200 column (GE Healthcare). Purified SE001, in 30 mM MES pH 6.5 and 300 mM sodium chloride, was aliquoted, flash frozen in liquid nitrogen, and stored at −80°C until purifications of the RXPF1 complex.

Single-chain relaxin-2 fused to an antibody IgG1 Fc fragment (SE301) was expressed as described above in Expi293F tetR cells. SE301 was purified using Protein G resin (GE Healthcare) in 20 mM HEPES pH 7.5, 150 mM sodium chloride. Purified SE301 was aliquoted, flash frozen in liquid nitrogen, and stored at −80°C until flow cytometry binding assays.

### Purification of the RXFP1–G_s_ complex

Purification of the RXFP1–G_s_ complex used a cell pellet from a 2 L induction of the RXFP1-miniG_s_399-20res Expi293F tetR stable cell line, and the purified proteins SE001 (single-chain relaxin-2), human Gβ_1γ2_, and Nb35-His-PrC. Gβ_1γ2_ and Nb35-His-PrC were expressed and purified according to previously published protocols^16^. The RXFP1-miniG_s_399-20res cell pellet was lysed through osmotic shock by stirring in 250 mL cold Lysis buffer containing 20 mM HEPES pH 7.5, 2 mM magnesium chloride, 1 μL benzonase (Sigma Aldrich), 1 protease inhibitor tablet (Thermo Fisher Scientific), and 100 nM SE001. Once stirring, iodoacetamide was added at a final concentration of 2 mg/mL. Lysed cells were centrifuged at 50,000 xg for 30 minutes at 4°C and the supernatant was decanted. Membranes were homogenized with a glass dounce (Thermo Fisher Scientific) in 270 mL Complexation buffer containing 20 mM HEPES pH 7.5, 350 mM sodium chloride, 20% (v/v) glycerol, 1 μL benzonase, and 1 protease inhibitor tablet. After homogenization, 960 μg Gβ_1γ2_, 740 μg Nb35-His-PrC (1:2:4 ratio of RXFP1:Gβ_1γ2_:Nb35-His-PrC), 100 nM SE001, and 0.2 U apyrase (New England Biolabs) were added to the membranes and the solution was stirred for 1 hour at 4°C. Next, 30 mL of 10X Detergent buffer [10% (w/v) lauryl maltose neopentyl glycol (L-MNG; Anatrace) and 1% (w/v) cholesterol hemisuccinate (CHS; Anatrace)] were added to the solution, dounce homogenization was repeated, and the solution was stirred for 2 hours at 4°C to extract the RXFP1–G_s_ complex from the membrane. After 2 hours, the solution was centrifuged at 50,000 xg for 30 minutes at 4°C and the supernatant was filtered with a glass fiber prefilter (Millipore). Calcium was added to the solubilized membranes at a final concentration of 2 mM and the solution was loaded by gravity flow over 3 mL M1 anti-FLAG resin (In house). This step of the purification was based on the N-terminal FLAG tag of RXFP1-miniG_s_399-20res. The M1 resin was equilibrated with Wash buffer 1 [0.1% (w/v) L-MNG, 0.01% (w/v) CHS, 350 mM sodium chloride, 20 mM HEPES pH 7.5, 2 mM calcium chloride, and 100 nM SE001]. After loading the solubilized membrane fraction, the column was subsequently washed with 50 mL each of Wash buffer 1, Wash buffer 2 [0.1% (w/v) L-MNG, 0.01% (w/v) CHS, 350 mM sodium chloride, 20 mM HEPES pH 7.5, 2 mM calcium chloride, 2 mM magnesium chloride, 5 mM adenosine 5’-triphosphate magnesium salt, and 100 nM SE001], and Wash buffer 3 [0.01% (w/v) L-MNG, 0.001% (w/v) CHS, 350 mM sodium chloride, 20 mM HEPES pH 7.5, 2 mM calcium chloride, and 100 nM SE001]. M1 resin was eluted with 0.01% (w/v) L-MNG, 0.001% (w/v) CHS, 350 mM sodium chloride, 20 mM HEPES pH 7.5, 0.5 mg/mL FLAG peptide (GenScript), and 100 nM SE001, then 2 mM calcium chloride was added to the eluted protein. Next, 0.5 mL anti-Protein C resin (In house) was equilibrated with Protein C wash buffer [0.005% (w/v) L-MNG,0.0005% (w/v) CHS, 350 mM sodium chloride, 20 mM HEPES pH 7.5, 2 mM calcium chloride, and 100 nM SE001]. The M1 elution with 2 mM calcium chloride was loaded onto the anti-Protein C resin by gravity flow, washed with 10 mL Protein C wash buffer, and eluted with 0.005% (w/v) L-MNG, 0.0005% (w/v) CHS, 350 mM sodium chloride, 20 mM HEPES pH 7.5, 0.5 mg/mL Protein C peptide (GenScript), and 100 nM SE001. This step of the purification was based on the Protein C tag of Nb35-His-PrC. The elution was concentrated with a 3 kDa molecular weight cutoff centrifugal concentrator (Millipore) and loaded onto a Sephadex S200 column (GE Healthcare) in SEC buffer [0.005% (w/v) L-MNG, 0.0005% (w/v) CHS, 350 mM sodium chloride, 20 mM HEPES pH 7.5, and 100 nM SE001]. Peak fractions from size exclusion were concentrated with a 3 kDa molecular weight cutoff centrifugal concentrator and either stored overnight at 4°C prior to grid freezing or immediately used to freeze cryo-EM grids.

### Cryo-EM grid preparation and data collection

Cryo-EM grids were prepared using QUANTIFOIL®holey carbon grids (400-mesh, copper, R1.2/1.3; Electron Microscopy Sciences). Grids were washed with ethyl acetate (Sigma Aldrich) and then glow-discharged at −20 mA for 60 seconds with a Pelco Easiglow. 3 μL of sample at 0.2 to 0.4 mg/mL was applied to the grids. Grids were plunge-frozen in liquid ethane using a Mark IV Vitrobot (Thermo Fisher Scientific) at 10°C and 100% humidity with a 10 second wait time, 3 to 4 second blot time, and blot force of 15.

Cryo-EM data for the RXFP1–G_s_ complex were collected in four separate sessions. Grids were imaged with a Titan Krios microscope (Thermo Fisher Scientific) at 300 kV with a Gatan BioQuantum GIF/K3 direct electron detection camera in counting mode. Movies were collected with 50 frames each at 81,000x, corresponding to 1.06 Å per pixel, and a total dose of around 52 electrons per Å^2^. Defocus values ranged from −0.8 to −2.3 μm. A total of 13,457 movies were collected across the four data collection sessions.

### Cryo-EM data processing

Cryo-EM data was collected in four separate session and initially processed individually, then particle stacks were merged to generate the final maps. Motion correction was carried out with MOTIONCOR2^49^ and CTF parameters were estimated with CTFFIND-4.1^50^. Particles were picked in RELION 3.1^51^ using Laplacian of Gaussian autopicking. A map from a previous small dataset collected on a Talos Arctica microscope and processed in RELION was used as an initial model. Particles were extracted with a box size large enough to include the entire complex (288 pixels). Multiple rounds of 2D classifications in RELION used a circular mask of 150 Å around only the micelle and G proteins as the first step of particle sorting for both 3D reconstructions. Masking of the micelle and G proteins was used to initially sort high quality particles from contamination because 2D classifications attempted for the entire particle showed weak signal for the ectodomain, likely indicating a region of higher flexibility. It became evident at this stage of data processing that the datasets showed a preferred orientation for one of the side views of the RXFP1–G_s_ complex. Processing 2D classifications of the preferred view separately from other views of the complex resulted in a larger number of particles from different orientations in the final particle stacks.

For the map of the 7TMs, masked 3D classifications of the 7TM domain bound to G proteins were performed in RELION. Following these classifications, iterative 3D refinements (using a mask of the micelle and G proteins) and Bayesian particle polishing steps were carried out in RELION. At this stage, particle stacks from different datasets were joined together, followed by additional 2D classifications with a 150 Å circular mask and masked 3D classifications. Finally, 3D focused classifications without alignments were carried out using masks for either the ECLs, TM helices, or helix8. These particle stacks were imported into cryoSPARC v3.1.0^39^ and used with Non-uniform Refinement (New). The cryoSPARC refined maps were post-processed in DeepEMhancer^52^ using the half-maps as input. This data processing workflow is described in **Fig. S2**.

For the map of the full-length RXFP1–G_s_ complex, rounds of unmasked 3D classifications were used in RELION after the 2D classifications described above. Next, the micelle and G proteins were subtracted from the particles and 3D focused classifications without alignments were carried out using a mask on the entire ectodomain of RXFP1. 3D refinements with a mask of the entire complex and iterative Bayesian particle polishing steps were then performed in RELION. Following particle polishing, particle stacks from different datasets were combined and 3D classifications with a mask around the entire complex were performed in RELION, followed by 3D refinement in RELION with a mask around the entire complex. Finally, local resolution estimation and filtering were performed in RELION on the final maps. This data processing workflow is described in **Fig. S3**. Continuous heterogeneity in the final particle stack was visualized using 3D variability analysis in cryoSPARC and shown in **Movie S1**.

### Model building and refinement

A combined focused map for the RXFP1 7TM domain bound to G proteins was generated using the DeepEMhancer post-processed maps as inputs in Phenix^53^. Model building for the RXFP1 7TM domain bound to heterotrimeric G_s_ and Nb35 was performed using DeepEMhancer focused and combined focused cryo-EM maps in *Coot*^54^. An initial model for the RXFP1 7TM domain (Gly395 to Gly709) was generated using trRosetta^55^ and manually fit into the DeepEMhancer focused cryo-EM map (for ECLs) in Chimera^56^. Intracellular and extracellular loops and the hinge region were removed from the trRosetta model and manually rebuilt in *Coot*. ECL2 residues Lys554-Tyr557 and hinge region residues Glu400-Leu403 were manually built with the coordinates of the human RXFP1 AlphaFold2 model^38^ overlayed in *Coot*. Initial models for miniG_s_399, Gβγ, and Nb35 were generated in MODELLER^57^ using coordinates from PDB ID 6GDG^58^. All models were refined with Phenix real-space refinement^53^ using secondary structure restraints against the DeepEMhancer combined focused map. Statistics for the final model were evaluated using MolProbity in a Phenix comprehensive validation (cryo-EM) job that used the map from cryoSPARC v3.1.0 Non-uniform Refinement (New) and the final model as inputs. Figures were prepared using PyMOL and ChimeraX^59^. Structural biology programs used in this work, other than cryoSPARC, were compiled and configured by SBGrid^60^. Refinement statistics are present in **Table S1** and representative images of cryo-EM map and model quality are shown in **Fig. S8**.

### Evolutionary coupling analysis

For comparing the constructed model to the strongest evolutionarily coupled pairs, we first downloaded the Uniprot canonical sequence for Q9HBX9 for residues 405-689 and then used the Jackhmmer software suite^61^ to build multiple sequence alignments across multiple bitscore thresholds based on the June 2019 download of the Uniref100 database^62^. We then chose an alignment with 352,511 sequences, with 90% of columns consisting of fewer than 30% gaps, to move forward with in analysis. We used the EVcouplings v0.0.5 software, available at <github.com/debbiemarkslab>, to identify evolutionary couplings (ECs) for this alignment.

To incorporate the LRR region, we ran the Q9HBX9 sequence for residues 120-757 in Jackhmmer, against the 02/2021 Uniref100 database for normalized bitscores between 0.1 and 0.9. We chose an alignment with 8.145 sequences with 86% of columns consisting of fewer than 30% gaps, and input it to v0.1.1 EVcouplings software. The strongest couplings were ranked based on assigning probabilities from a logistic regression model new to v0.1.1. This version of the pipeline was also used for plotting all contact maps.

### Crosslinking mass spectrometry

The RXFP1–G_s_ complex used for CLMS was prepared as described above and crosslinking reactions were carried out the following day, after storage overnight at 4°C. CLMS was performed as previously described^41^. Briefly, crosslinking reactions were carried out for 1 h at room temperature in 100 mM MOPS Buffer, pH 6.5 with 50 mM EDC, ~24 mM EDDA linker, and 20 mM sulfo-NHS. Reactions were quenched with hydroxylamine to a final concentration of 100 mM. Samples were reduced for 1 h in 2% SDS and 5 mM TCEP followed by alkylation with 10 mM iodoacetamide in the dark for 30 min and quenching with 5 mM DTT for 15 min. Samples were then processed with the SP3^63^ method and digested with trypsin (Promega) at 1:25 enzyme:substrate ratio overnight at 37°C. Digested peptides were acidified with 10% formic acid to pH ~2 and desalted using stage tips with Empore C18 SPE Extraction Disks (3M) and dried under vacuum.

Sample was reconstituted in 5% formic acid (FA)/5% acetonitrile and analyzed in the Orbitrap Eclipse Mass Spectrometer (Thermo Fischer Scientific) coupled to an EASY-nLC 1200 (Thermo Fisher Scientific) ultra-high pressure liquid chromatography (UHPLC) pump, as well as a high-Field Asymmetric waveform Ion Mobility Spectrometry (FAIMS) interface. Peptides were separated on an in-house packed 100 μm inner diameter column packed with 35 cm of Accucore C18 resin (2.6 um, 150 Å, ThermoFisher), using a gradient consisting of 5–35% (ACN, 0.125% FA) over 135 min at ~500 nL/min. The instrument was operated in data-dependent mode. FTMS1 spectra were collected at a resolution of 120K, with an automated gain control (AGC) target of 5 × 10^5^, and a max injection time of 50 ms. The most intense ions were selected for MS/MS for 1s in top-speed mode, while switching among three FAIMS compensation voltages (CV): −40, −60, and −80 V in the same method. Precursors were filtered according to charge state (allowed 3 <= z <=7), and monoisotopic peak assignment was turned on. Previously interrogated precursors were excluded using a dynamic exclusion window (60 s ± 7 ppm). MS2 precursors were isolated with a quadrupole mass filter set to a width of 0.7 m/z and analyzed by FTMS2, with the Orbitrap operating at 30K resolution, an AGC target of 100K, and a maximum injection time of 150 ms. Precursors were then fragmented by high-energy collision dissociation (HCD) at a 30% normalized collision energy.

Mass spectra were processed and searched using the PIXL search engine^41^. The sequence database contained proteins identified at 1% FDR in non-cross-linked Comet^64^ search. For PIXL search, precursor tolerance was set to 15 ppm and fragment ion tolerance to 10 ppm. Methionine oxidation was set as a variable modification in addition to mono-linked mass of +130.110613 for EDDA. Crosslinked peptides were searched assuming zero-length (−18.010565) and EDDA crosslinker +112.100048. Crosslinked searches considered 60 protein sequences to ensure sufficient statistics for FDR estimation. Matches were filtered to 1% FDR on the unique peptide level using linear discriminant features as previously described^41^.

### Docking

HADDOCK^44^ was used to dock the relaxin-2–LRR interaction. Docking used the human relaxin-2 X-ray crystal structure (PDB ID: 6RLX)^65^ and a model of the LRRs from residues 104-391 of the AlphaFold2^38^ prediction for human RXFP1. Residues from CLMS studies that were part of crosslinks between relaxin-2 and RXFP1 were used as active restraints in the docking run. These CLMS residues were Glu14^B-chain^ of relaxin-2, and Glu206, Glu299, and Glu351 of RXFP1. Residues identified to be important for relaxin-2 binding from published mutations in radioligand binding assays were also used as active restraints^32,42,43^. These residues included Arg13^B-chain^, Arg17^B-chain^, and Ile20^B-chain^ of relaxin-2 and Trp202, Ile204, Leu226, Asp253, Glu255, Glu299, and Asp301 of RXFP1. The resulting docking models were analyzed according to the HADDOCK scoring function and fit into the low resolution cryo-EM map of the RXFP1 ectodomain in ChimeraX^59^.

### Flow cytometry binding assay

A flow cytometry assay was used to measure the binding of an Fc-tagged relaxin-2 protein, SE301^17^, to Expi293F cells transfected with human RXFP1 or empty pcDNA-Zeo-tetO vector. Expi293F tetR cells were grown in Expi293 media and transfected using FectoPRO (Polyplus), according to the manufacturer’s protocols. The cells were enhanced 24 hours post-transfection with 0.4% glucose and induced 48 hours post-transfection with 4 μg/mL doxycycline and 5 mM sodium butyrate. After 24 hours of induction, cells were harvested by spinning at 200 xg for 5 minutes at 4°C and washed once with HBS with 1% (v/v) FBS and 2 mM calcium chloride (Buffer B). Cells were plated into a V-bottom 96-well plate (Corning) at 100,000 cells/well and blocked by incubation in Buffer B for 30 minutes at 4°C. After blocking, cells were centrifuged at 200 xg for 5 minutes at 4°C, resuspended in 100 μL of Buffer B containing 500 nM SE301, and incubated for 1 hour at 4°C. Cells were then centrifuged at 200 xg for 5 minutes at 4°C, washed twice with 200 μL Buffer B, and resuspended in 100 μL Buffer B containing 100 nM M1 anti-FLAG antibody labeled with Alexa Fluor 488 (In house) and Alexa Fluor 647 anti-human IgG Fc (BioLegend) diluted 1:100 (v/v). Cells were incubated in secondary antibodies for 30 minutes at 4°C, washed once with 200 μL Buffer B, and resuspended in 100 μL Buffer B for flow cytometry.

Samples were analyzed on a CytoFLEX flow cytometer (Beckman Coulter) and gated according to plots of FSC-A/SCA-A, FSC-A/FSC-H, and receptor expression according to Alexa Fluor 488 M1 anti-FLAG antibody binding (**Fig. S11**). The receptor expression gate was drawn by comparing empty vector and wild type RXFP1-transfected cells. Approximately 500 events/sample were collected from cells expressing receptor for human RXFP1-transfected cells or post-FSC-A/FSC-H gating for empty vector-transfected cells. The data were plotted and analyzed in FlowJo and GraphPad Prism. For the binding assay in **Fig. 4**, the E206A and wild type RXFP1 samples expressed very similarly, so cell surface expression and SE301 binding were plotted separately. For comparing the binding of multiple constructs in **Fig. S5**, ratios of SE301 binding to receptor expression were calculated in order to normalize for the differences in RXFP1 construct expression levels.

A flow cytometry competition binding assay was used to measure SE001 binding to human RXFP1-expressing Expi293F cells. Expi293F cells were transfected, harvested, and blocked as stated above. After blocking in Buffer B, cells were incubated with 200 nM SE301 (Fc-tag) and increasing concentrations of SE001 (His-tag). After 1 hour of incubation at 4°C, the reaction was terminated by centrifugation at 200 x g for 5 minutes at 4°C and cells were washed twice with 200 μL Buffer B. In order to detect SE301 and measure receptor expression, cells were then stained with Alexa Fluor 488 M1 anti-FLAG and Alexa Fluor 647 anti-human IgG Fc and analyzed by flow cytometry as stated above. Data points were calculated as a percentage of wild type RXFP1 SE301 binding and plotted in GraphPad Prism (**Fig. S1f**).

### Molecular dynamics

The initial model was built from the cryo-EM structure reported here. The missing segments in ECL2, ICL3 and TM6 were generated by MODELLER v9.15^66^. PACKMOL-Memgen^67^ was used to assign the side-chain protonation states and embed the models in a bilayer of POPC lipids. The systems were solvated in a periodic box of explicit water and neutralized with 0.15 M of Na^+^ and Cl^-^ ions. We used the Amber ff14SB^68^ and lipid 14^69^ force fields, the TIP3P water model^70^ and the Joung-Cheatham ion parameters^71^. For the simulations of 7TM deactivation, a Na^+^ ion was placed at the conserved Na^+^-binding site (between Asp451^2.50^ and Ser495^3.35^). For the simulations of autoactivation, the truncated AlphaFold2^38^ model was used to build the truncated 7TM and ^half^LRRs-7TM forms in which Asp451^2.50^ was protonated.

After energy minimization, all-atom MD simulations were carried out using Gromacs 5.1^72^ patched with the PLUMED 2.3 plugin^73^. The LINCS algorithm^74^ was applied to constrain bonds involving hydrogen atoms, allowing for a time step of 2 fs. Each system was gradually heated to 310 K and pre-equilibrated during 10 ns of brute-force MD in the *NPT-*ensemble. The replica exchange with solute scaling (REST2)^75^ technique was used to enhance the conformational sampling. A total of 64 replicas of simulations were performed in the *NVT* ensemble. REST2 is a type of Hamiltonian replica exchange simulation scheme. Besides the original simulation, many replicas of the same system were simulated simultaneously. The additional replicas have modified free energy surfaces, in which the energy barriers are easier to cross than in the original simulation system. By frequently swapping the replicas and the original system during the MD, the simulations “travel” on different free energy surfaces and easily visit various conformational zones. Finally, only the samples on the original free energy surface are collected. The additional replicas are artificial to ease barrier crossing, which are discarded after the simulations. REST2, in particular, modifies the free energy surfaces by scaling (reducing) the force constants of the “solute” molecules in the simulation system. In this case, the protein was considered as “solute”–the force constants of its van der Waals, electrostatic and dihedral terms were subject to scaling–in order to facilitate the conformational changes. The scaling factors were generated using the Patriksson-van der Spoel approach^76^ and effective temperatures ranging from 310 K to 1000 K. Exchange between replicas was attempted every 1000 simulation steps. This setup resulted in an average exchange probability of ~40% for the 7TM and ~20% for the ^half^LRRs-7TM systems, respectively. We performed 80 ns × 64 replicas of REST2 MD in the *NVT* ensemble for each system. The first 30 ns were discarded for equilibration. The simulation trajectories on the original unmodified free energy surface was reassembled and analyzed.

