## Supplementary_information for "The relaxin receptor RXFP1 signals through a mechanism of autoinhibition"

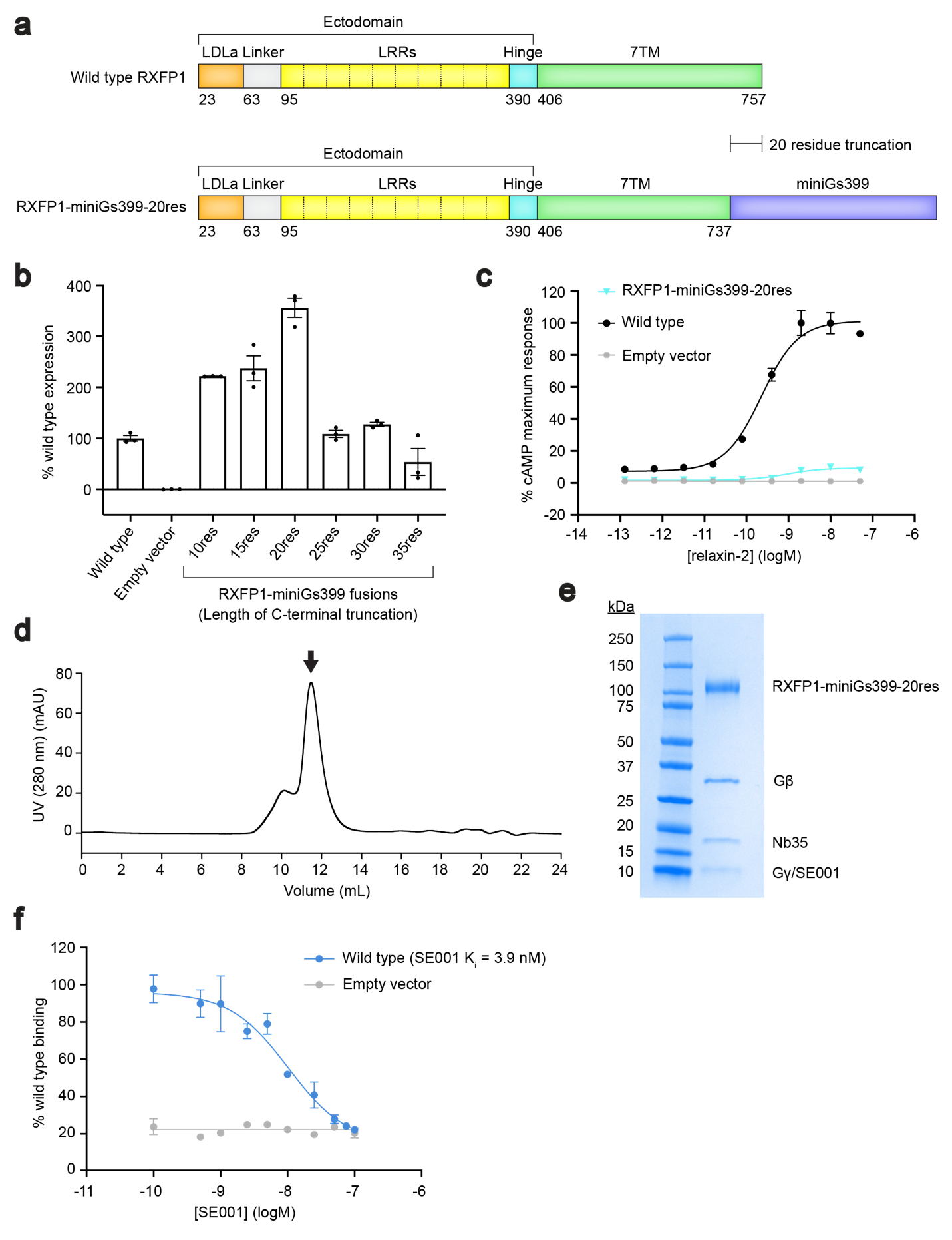

**Fig. S1. Engineering and purification of the RXFP1–G_s_ complex. a**, Diagram of the primary structure of RXFP1 domains versus the RXFP1-miniG_s_399-20res fusion construct. **b**, Flow cytometry cell surface expression tests in Expi293F tetR cells for RXFP1-miniG_s_ fusion constructs. Data is mean ± s.e.m., n=3 technical replicates. **c**, G_s_ signaling assay comparing the signaling levels of wild type RXFP1 versus RXFP1-miniG_s_399-20res in response to relaxin-2. **d**, Size exclusion chromatography profile for the RXFP1–G_s_ complex. Arrow indicates the peak fractions pooled for RXFP1–G_s_. **e**, Coomassie-stained SDS-PAGE gel for the RXFP1–G_s_ complex. **f,** Flow cytometry competition binding assay for SE001^1^. SE001 competes with 200 nM SE301 for binding to wild type RXFP1. The K_i_ for SE001 was calculated to be 3.9 nM; data is mean ± s.e.m., n=3 technical replicates.

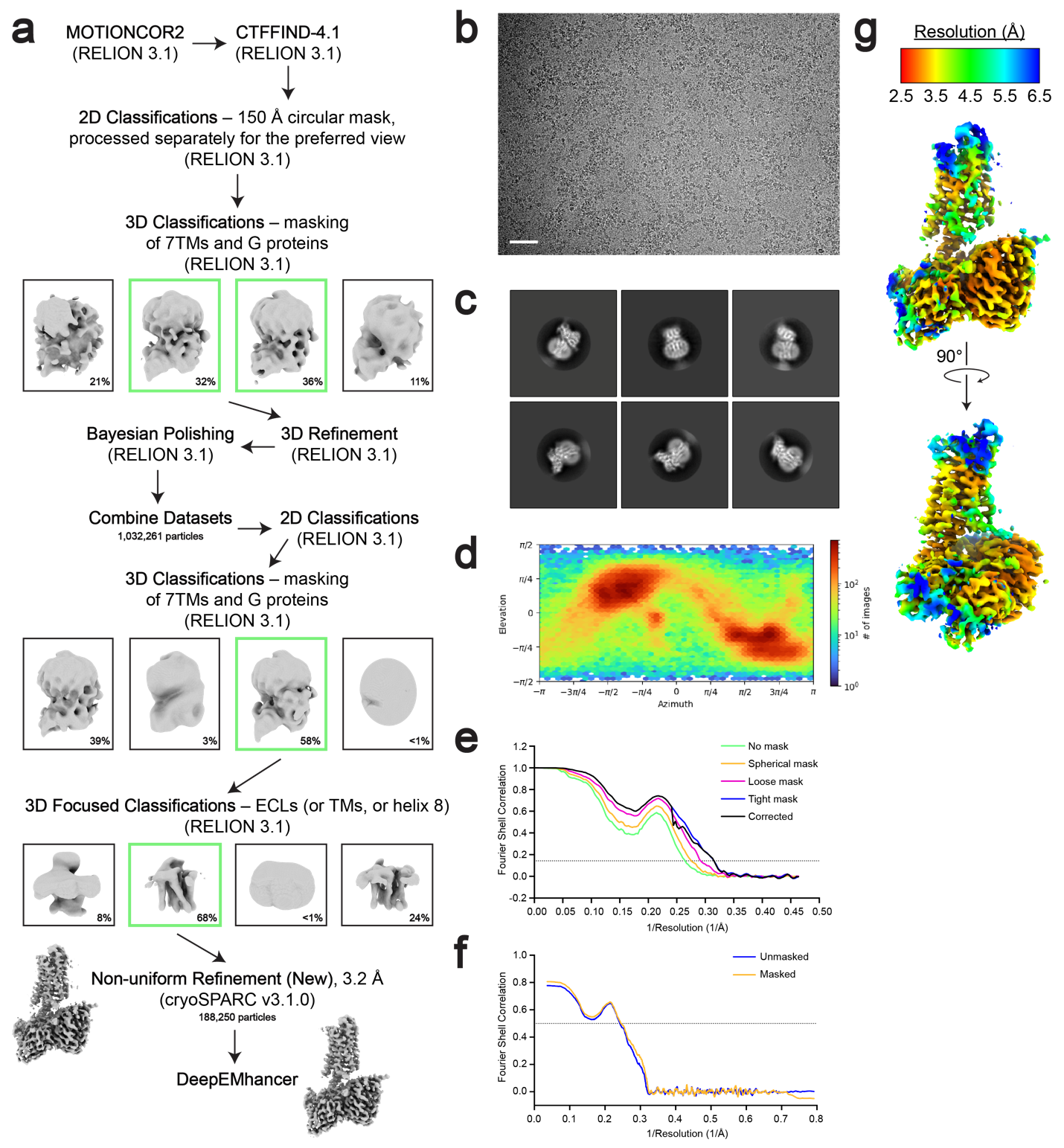

**Fig. S2. Cryo-EM data processing for the 7TM domain of RXFP1–G_s_. a**, Cryo-EM data processing scheme for the 7TM domain of RXFP1 in complex with G_s_. Shown are representative processing steps for one of four individual datasets and the steps used for the combined datasets. **b**, Representative micrograph from the RXFP1–G_s_ complex datasets (Scale bar = 50 nm). **c**, Two-dimensional class averages for the 7TM domain of RXFP1 and G proteins. **d**, Angular distribution of particles in the final refinement for the 7TM domain with G proteins. **e**, Fourier shell correlation (FSC) used to determine the overall map resolution. **f**, Map to model FSC curve. **g**, cryoSPARC non-uniform refinement map colored by local resolution.

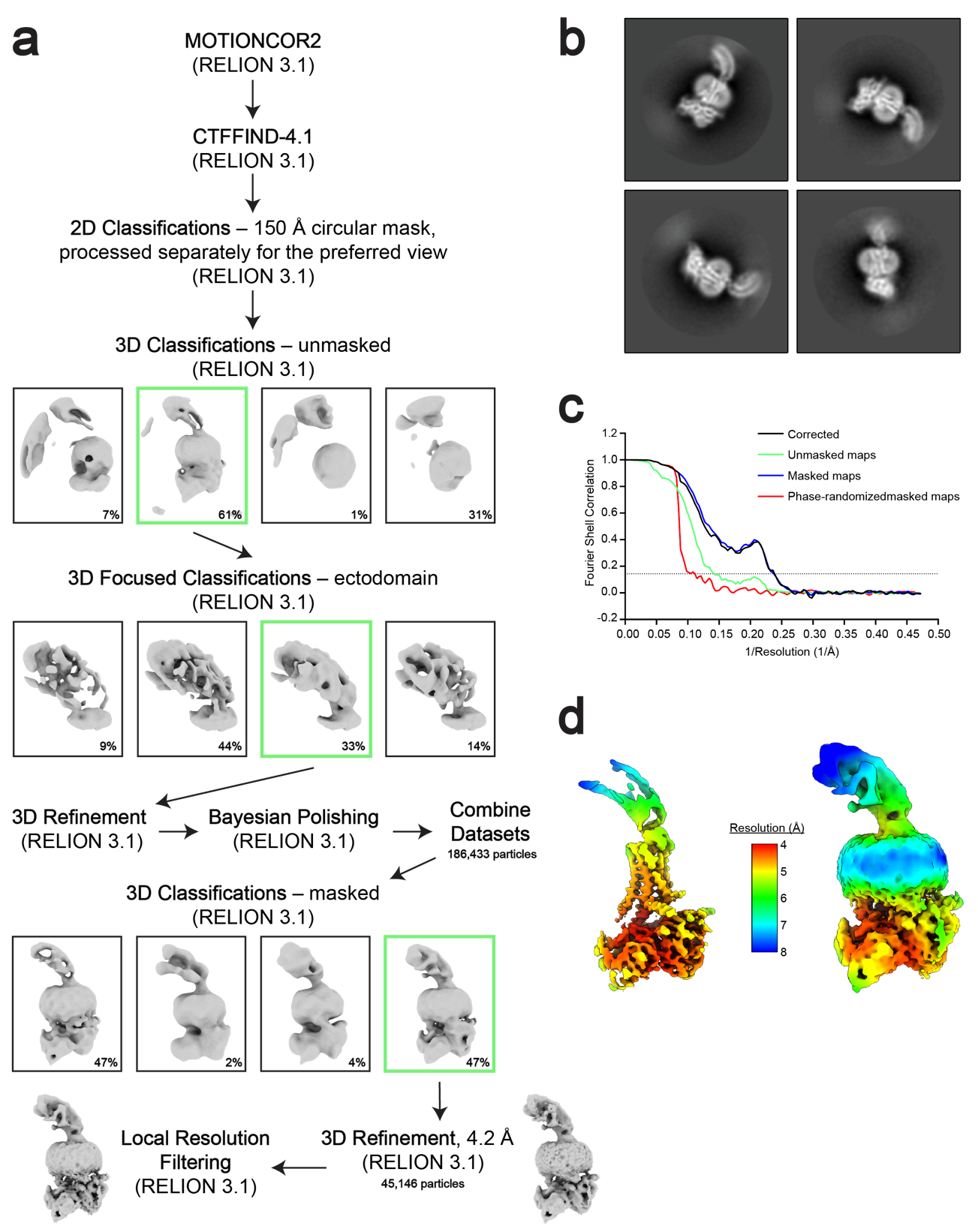

**Fig. S3. a**, Cryo-EM data processing scheme for the full-length RXFP1–G_s_ complex. Shown are representative processing steps for one of four individual datasets and the steps used for the combined datasets. **b**, Two-dimensional class averages for the full-length RXFP1–G_s_ complex. **c**, FSC used to determine the overall resolution of the map. **d,** RELION map of the full-length RXFP1 complex colored by local resolution at two different contour levels to display both the TM helices and receptor ectodomain.

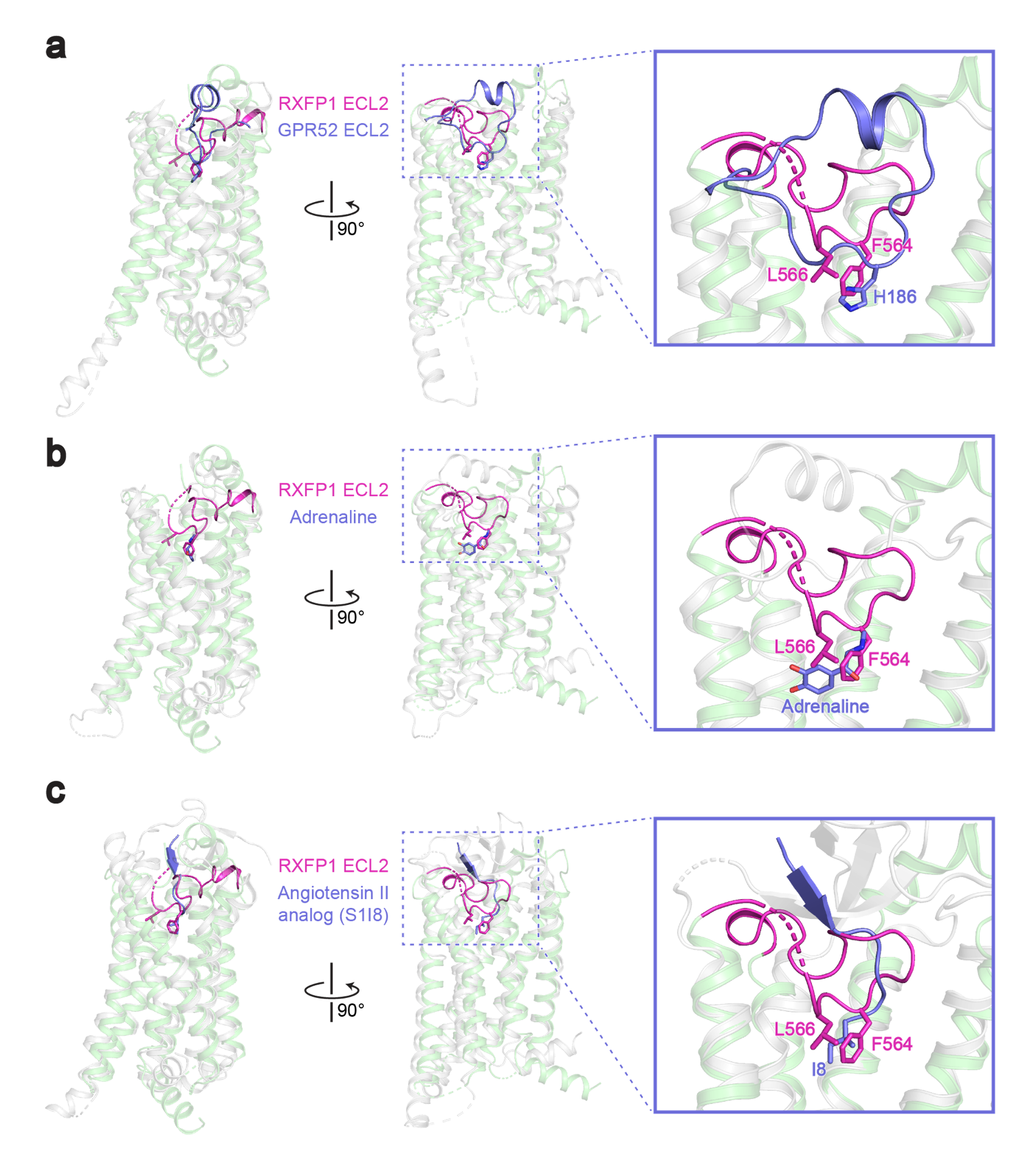

**Fig. S4. Alignments of RXFP1’s ECL2 with GPR52 and family A orthosteric agonists. a-c,** Alignment of active-state RXFP1 (green, with ECL2 in magenta) with GPR52 (gray with ECL2 in purple; PDB ID: 6LI3)^2^ (**a**), the β_2_ adrenergic receptor (gray) bound to adrenaline (purple; PDB ID: 4LDO)^3^ (**b**), and the angiotensin II type I receptor (gray) bound to the angiotensin II analog S1I8 (purple; PDB ID: 6DO1)^4^ (**c**).

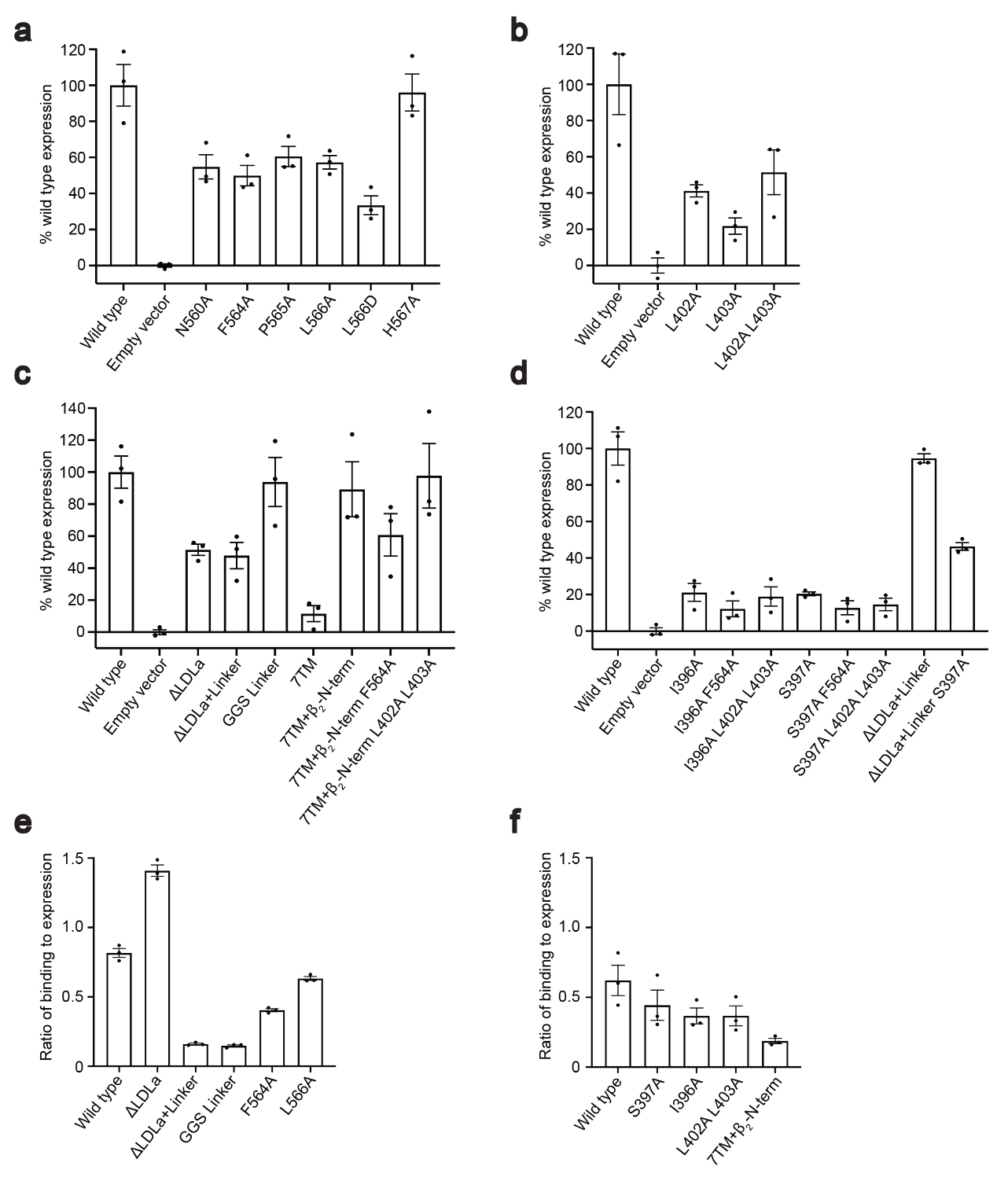

**Fig. S5. Cell surface expression and SE301 binding for RXFP1 constructs. a-d,** Flow cytometry cell surface expression tests with HEK293T cells for RXFP1 ECL2 mutants (**a**), Leu402 and Leu403 hinge region mutants (**b**), ectodomain truncation constructs (**c**), and evolutionary coupling analysis Ile396 and Ser397 hinge mutants (**d**). Data is mean ± s.e.m., n=3 technical replicates. **e-f,** Ratio of SE301 (Fc-tagged relaxin-2) binding to receptor expression for flow cytometry binding assays in Expi293F cells^1^. Data is mean ± s.e.m., n=3 technical replicates. Deletion or mutations to the linker region reduce the ratio of binding to expression, while LDLa deletions and ECL2 mutations retain an ability to bind SE301 (**e**). Ectodomain deletions (7TM + β_2_N-term) reduce the ratio of binding to expression, while mutations to the hinge region maintain an ability to bind SE301 (**f**).

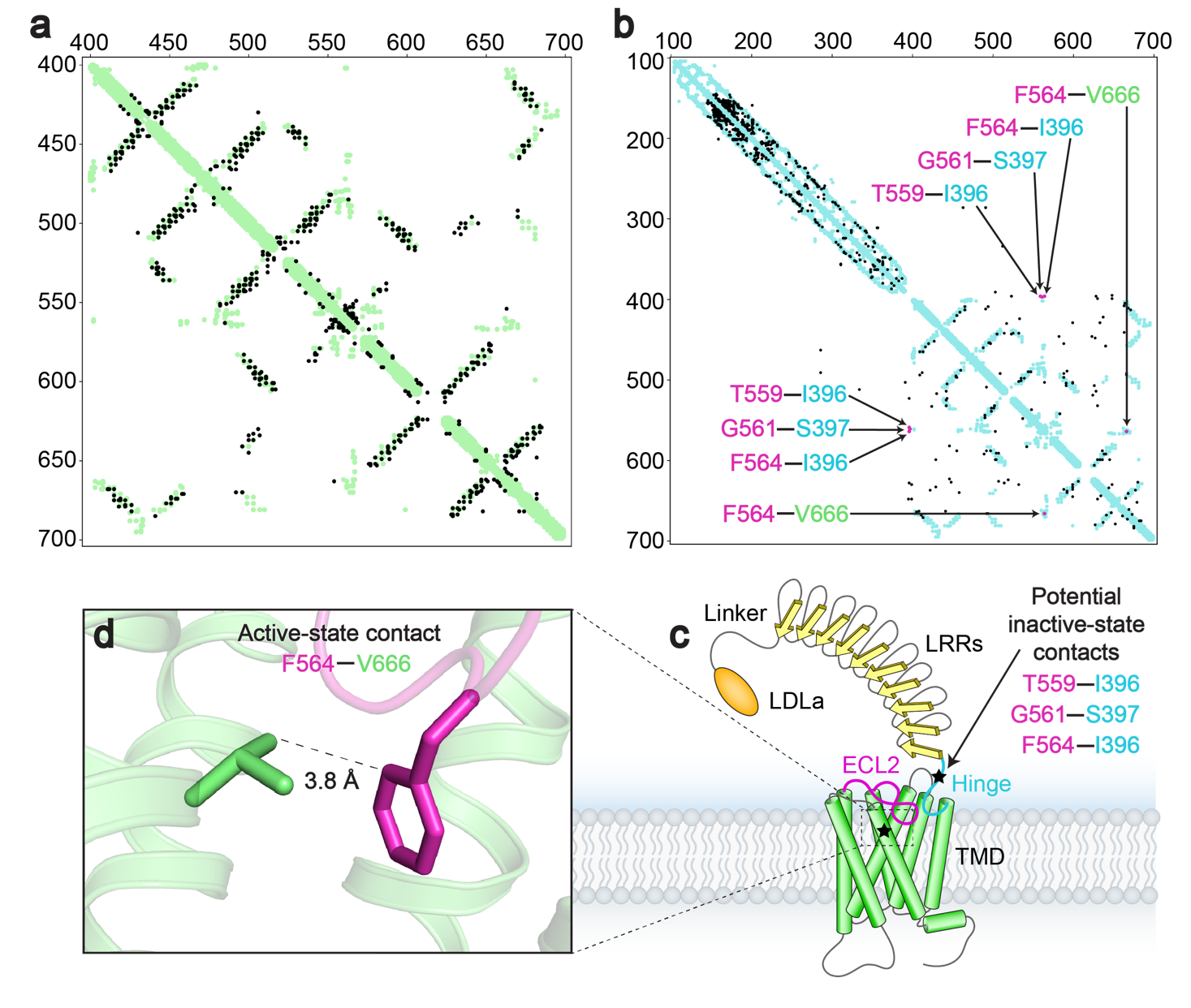

**Fig. S6. a**, Evolutionary couplings for RXFP1 residues 405-689 (black) compared to the active-state structure contacts (green) show close agreement between predicted contacts from ECs and the cryo-EM model. **b**, Evolutionary couplings for RXFP1 residues 120-757 (black) compared to the active-state 7TM structure and LRR AlphaFold2^5^ model contacts (blue), highlighting ECL2 evolutionary couplings that provide insight into two potential loop conformations in magenta (T559^ECL2^–Ile396, Gly561^ECL2^–Ser397, Phe564^ECL2^–Ile396, Phe564^ECL2^–Val666^7.38^). **c**, Diagram of RXFP1 domains. Stars indicate two regions of ECL2 predicted contacts from ECs, TM7 and the hinge region. **d**, The Phe564^ECL2^ and Val666^7.38^ residues from evolutionary coupling analysis are in close contact in the RXFP1 active-state structure.

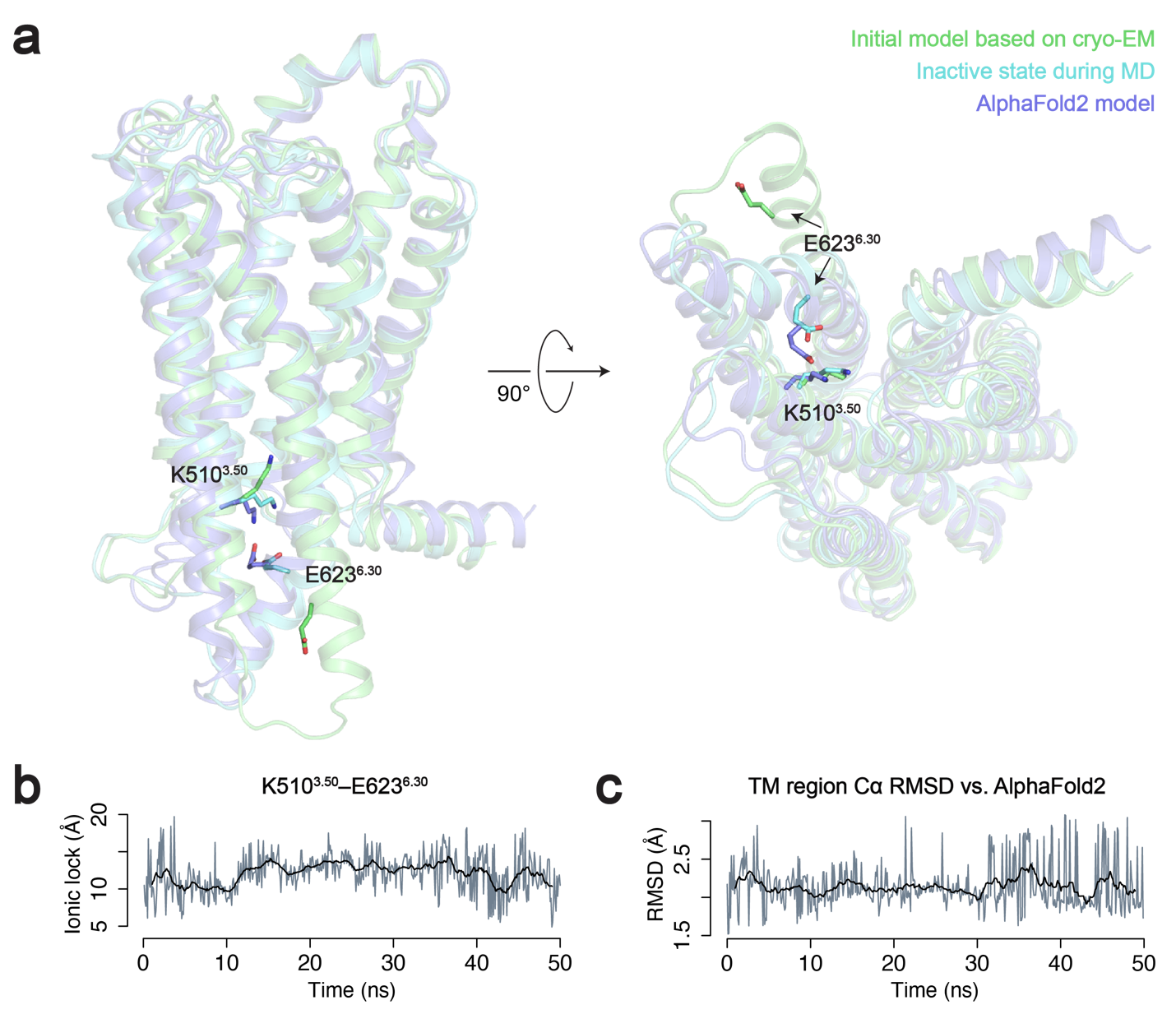

**Fig. S7. a,** The truncated 7TM domain is deactivated by adding a sodium in the conserved sodium-binding site. This leads to an inactive state during the simulations that closely resembles the AlphaFold2 model. **b-c,** Ionic lock distance and Cα RMSD of the TM region with respect to the AlphaFold2 model.

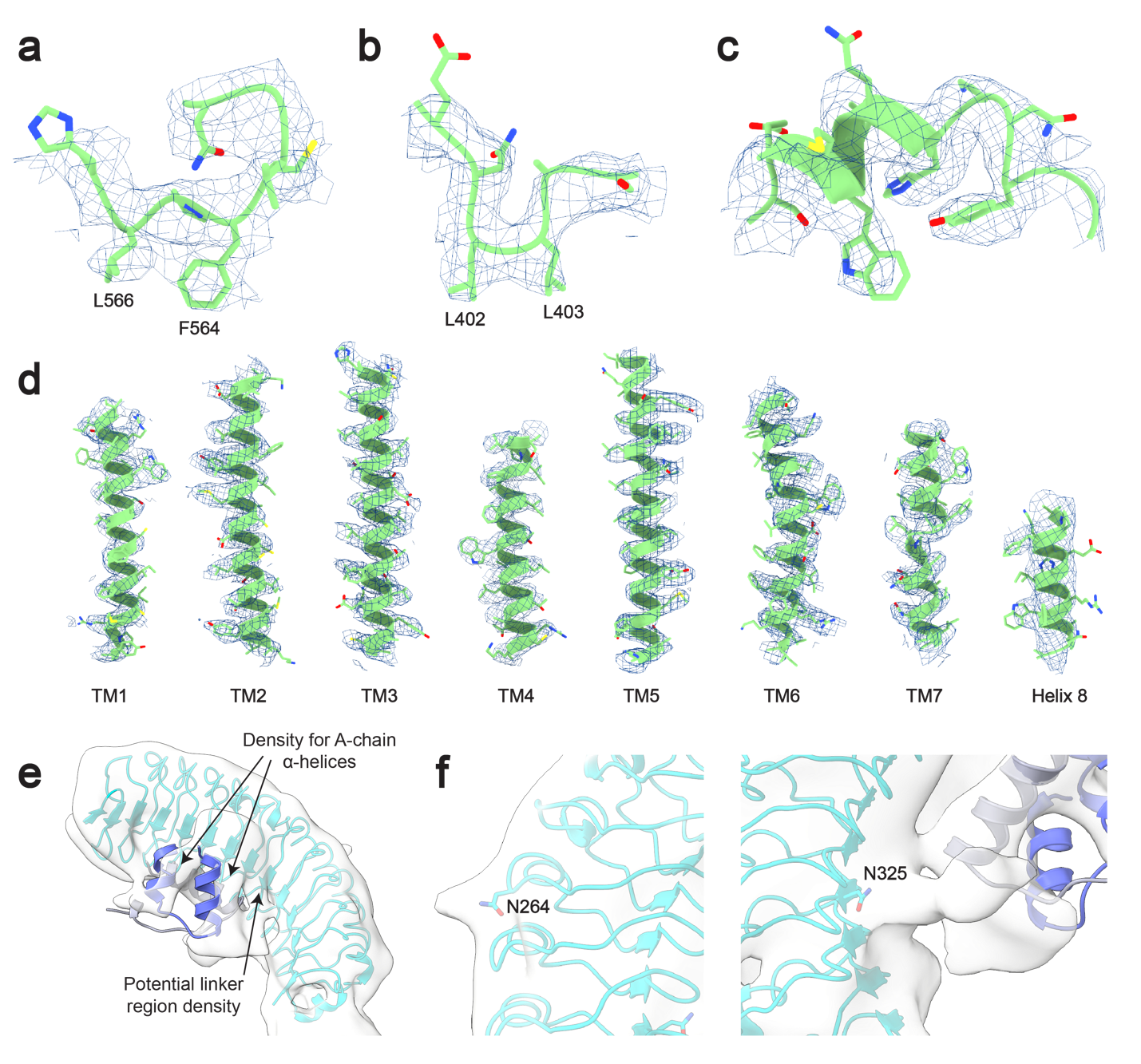

**Fig. S8. a-c,** Cryo-EM map and model for ECL2 (**a**), the hinge region (**b**), and ECL1 (**c**). **d**, Cryo-EM map and models for TMs 1-7 and Helix 8. **e**, Potential linker region density next to the relaxin-2 A-chain helices in the low-resolution ectodomain cryo-EM map. **f**, Low-resolution features in the ectodomain cryo-EM map at predicted sites of N-linked glycosylation.

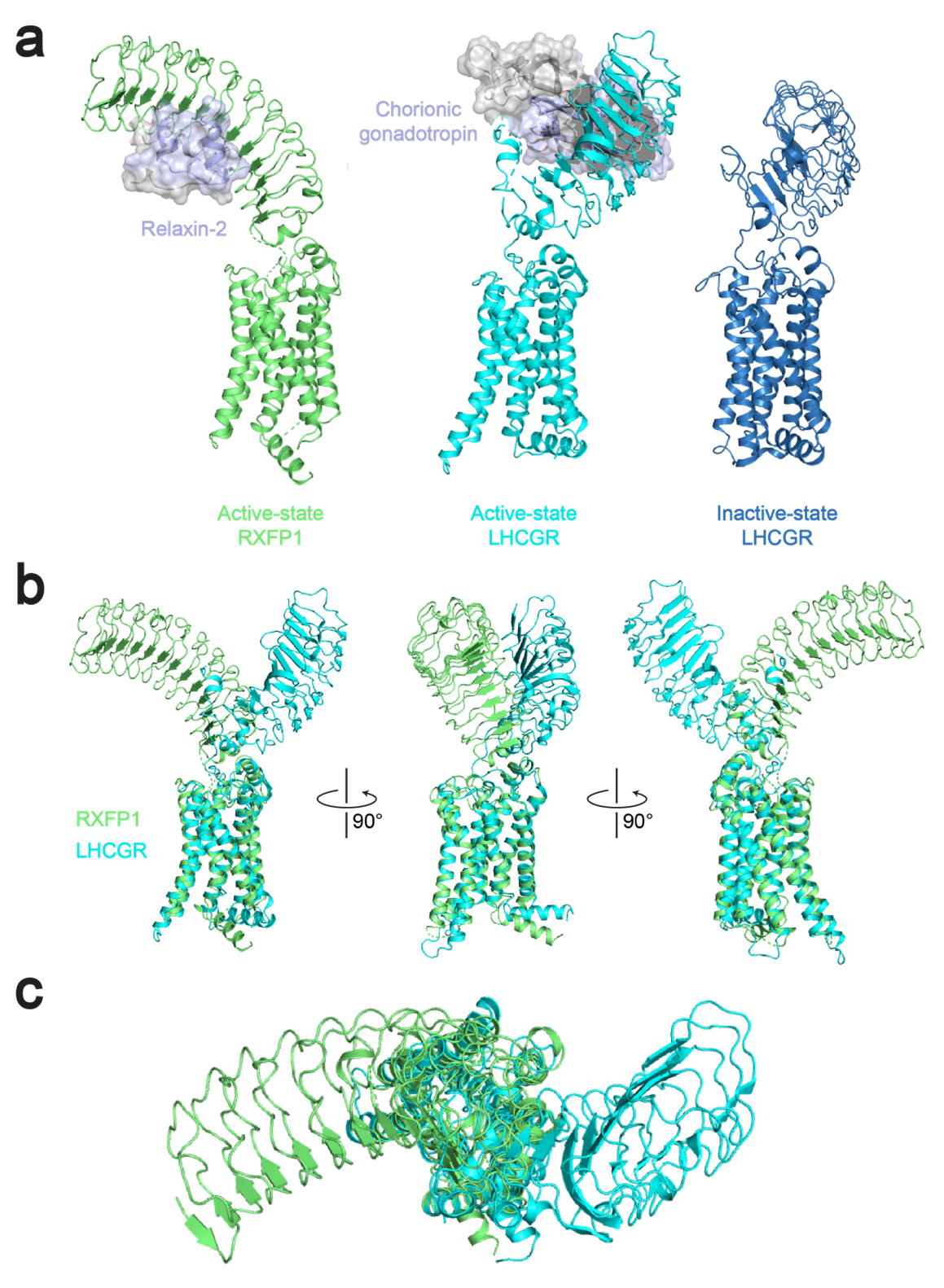

**Figure S9. Comparison of LHCGR and RXFP1 structures. a-b,** The hybrid model of active-state RXFP1 (based on our 7TM domain structure, cryo-EM maps, and AlphaFold2 model of the LRRs with docked relaxin-2 hormone) compared to active-state LHCGR (PDB ID: 7FIG) and inactive-state LHCGR (PDB ID: 7FIJ)^6^. The receptors are aligned on the 7TM domain. **b-c,** Side views (**b**) and top view (**c**) of the hybrid model of active-state RXFP1 compared to active-state LHCGR (PDB ID: 7FIG). The receptors are aligned on the 7TM domain and the ligands (relaxin-2 and chorionic gonadotropin) not displayed in order to highlight differences in active-state LRR orientations.

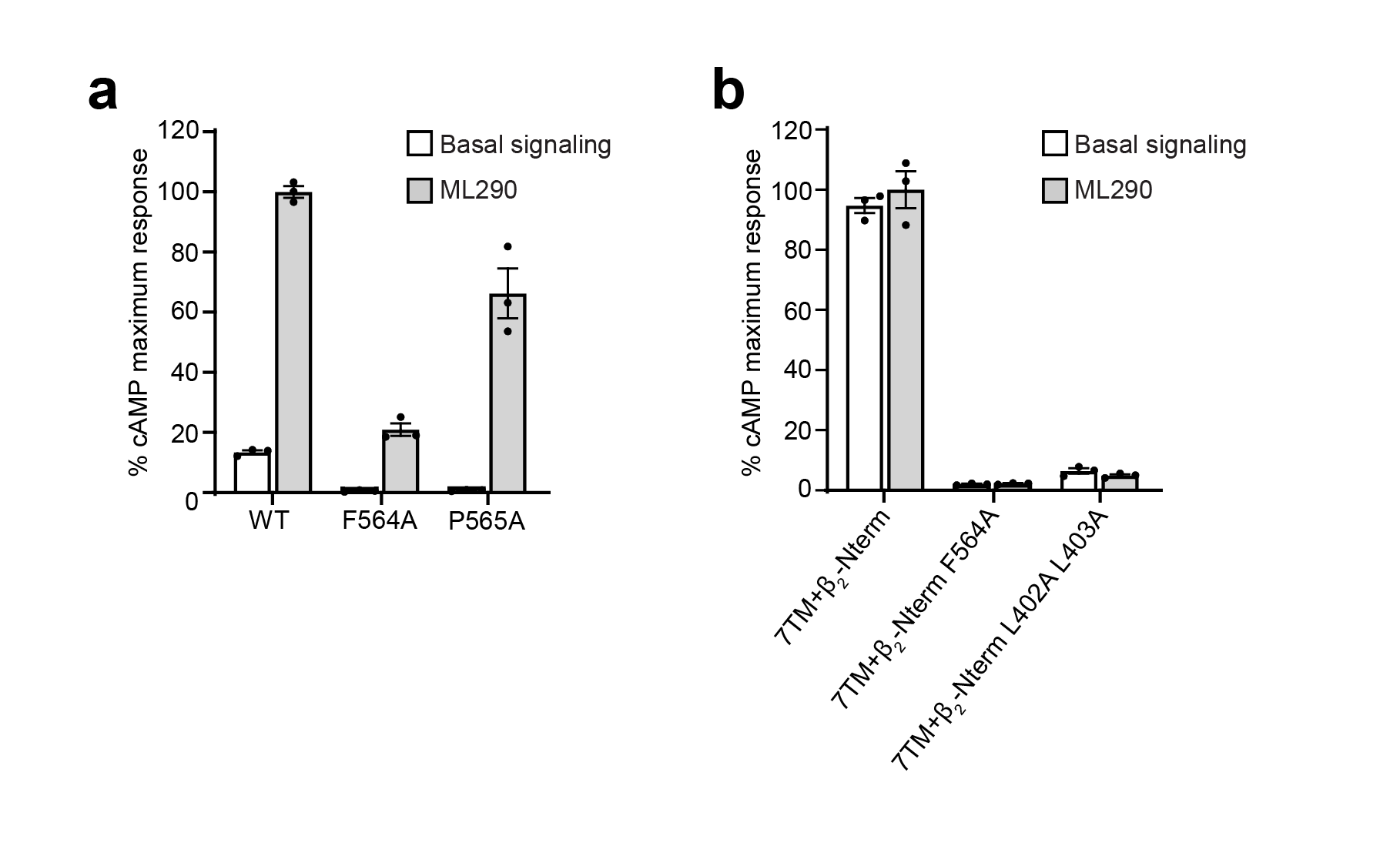

**Figure S10. Signaling of RXFP1 constructs by the small molecule agonist ML290. a,** G_s_ signaling at wild type RXFP1 and ECL2 mutants at basal levels and in response to 490 nM ML290. **b,** G_s_ signaling at constructs of the RXFP1 7TM domain fused to the β_2_ adrenergic receptor N-terminus at basal levels and in response to 490 nM ML290.

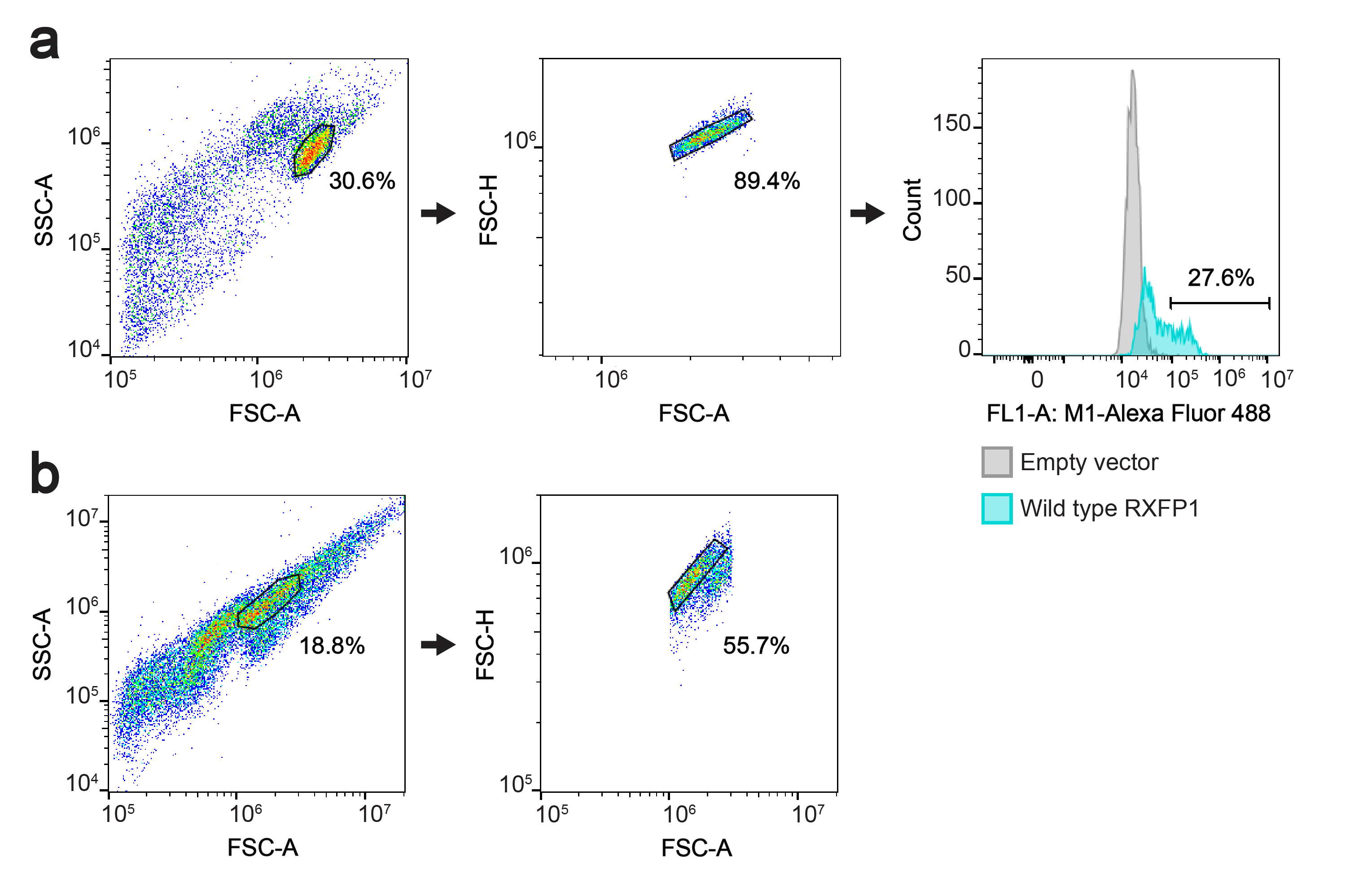

**Figure S11. Representative flow cytometry gating strategies. a-b,** Flow cytometry plots illustrating the gating strategies used for the flow cytometry binding assay with Expi293F cells in **Fig. 4** and **Fig. S5e,f** (**a**) and cell surface expression tests with HEK293T cells in **Fig. S5a-d** (**b**).

**Movie S1, related to Fig. 4 and Fig. S3. Dynamics of RXFP1’s active-state ectodomain.** Continuous heterogeneity within the final cryo-EM particle stack used for refinement of the full-length RXFP1–G_s_ complex. Flexibility of the ectodomain with respect to the 7TM domain and G proteins was visualized using 3D variability analysis in cryoSPARC.

**Table S1. Cryo-EM data collection, refinement, and validation statistics.**

Abbreviations: 7TM, RXFP1 masking the 7TM domain and G proteins; FL, full-length RXFP1

|  | RXFP1–G_s_-7TM | RXFP1–G_s_-FL |
| --- | --- | --- |
|  | (PDB 7TMW) (EMDB-26003) | (EMDB-26004) |
| Cryo-EM data collection and processing |  |  |
| Magnification | 81,000 | 81,000 |
| Voltage (kV) | 300 | 300 |
| Electron exposure (e-/ Å^2^) | ~52 | ~52 |
| Defocus range (μm) | -0.8 to -2.3 | -0.8 to -2.3 |
| Pixel size (Å) | 1.06 | 1.06 |
| Symmetry | C1 | C1 |
| Initial particle images (no.) | 15,826,542 | 15,826,542 |
| Final particle images (no.) | 188,250 | 45,146 |
| Map resolution (Å) | 3.2 | 4.2 |
| FSC threshold | (0.143) | (0.143) |
| Model refinement and validation |  |  |
| Initial model used (PDB) | Model generated from 6GDG  chains B, C, D, and E | |
| Model resolution (Å) | 4.1 |  |
| FSC threshold | (0.5) |  |
| Map sharpening *B* factor | DeepEMhancer |  |
| Model composition |  |  |
| Non-hydrogen atoms | 7320 |  |
| Protein residues | 930 |  |
| Ligands | 0 |  |
| R.m.s. deviations |  |  |
| Bond lengths (Å) | 0.003 |  |
| Bond angles (Å) | 0.701 |  |
| Validation |  |  |
| MolProbity score | 1.62 |  |
| Clashscore | 9.24 |  |
| Poor rotamers (%) | 0.00 |  |
| Ramachandran plot |  |  |
| Favored (%) | 97.35 |  |
| Allowed (%) | 2.65 |  |
| Disallowed (%) | 0.00 |  |

**Table S2. G_s_ signaling and expression data for RXFP1 constructs in Fig. 2c,d and Fig. S5a,b.**

^†^Mean ± s.e.m., n=3 technical replicates.
ND, not determined.

| Construct | pEC_50_ | E_max_ (%) | Cell surface expression (%)^†^ |
| --- | --- | --- | --- |
| Wild type | 9.8 ± 0.1 | 100 ± 2.3 | 100 ± 12 |
| Empty vector | ND | 1 ± 0.1 | 0 ± 1 |
| N560A | 9.3 ± 0.1 | 92 ± 2.6 | 55 ± 7 |
| F564A | 8.4 ± 0.1 | 16 ± 0.8 | 50 ± 6 |
| P565A | 8.6 ± 0.1 | 65 ± 3.6 | 61 ± 6 |
| L566A | 9.0 ± 0.1 | 77 ± 2.5 | 57 ± 4 |
| L566D | 8.0 ± 0.1 | 11 ± 0.4 | 33 ± 5 |
| H567A | 9.0 ± 0.05 | 96 ± 2.3 | 96 ± 10 |
| Wild type | 9.8 ± 0.1 | 100 ± 3.3 | 100 ± 17 |
| Empty vector | ND | 2 ± 0.1 | 0 ± 4 |
| L402A | 8.8 ± 0.1 | 32 ± 1.6 | 41 ± 3 |
| L403A | 8.8 ± 0.1 | 23 ± 0.7 | 22 ± 5 |
| L402A L403A | ND | 1 ± 0.05 | 51 ± 12 |

**Table S3. G_s_ signaling and expression data for RXFP1 constructs in Fig. 2e,f and Fig. S5c,d.**

^†^Mean ± s.e.m., n=3 technical replicates.

^‡^Mean ± s.e.m., n=9 technical replicates.

| Construct | Basal signaling (%)^‡^ | Relaxin-2 signaling (%)^‡^ | Cell surface expression (%)^†^ |
| --- | --- | --- | --- |
| Wild type | 8 ± 0.6 | 100 ± 5.3 | 100 ± 10 |
| Empty vector | 2 ± 0.2 | 2 ± 0.1 | 0 ± 2 |
| ΔLDLa | 8 ± 0.5 | 9 ± 0.6 | 51 ± 3 |
| ΔLDLa+Linker | 10 ± 0.6 | 10 ± 0.7 | 48 ± 8 |
| GGS Linker | 12 ± 0.6 | 12 ± 0.7 | 94 ± 15 |
| 7TM | 11 ± 0.5 | 10 ± 0.5 | 12 ± 5 |
| 7TM+β_2_-Nterm | 72 ± 3.2 | 70 ± 3.1 | 89 ± 17 |
| 7TM+β_2_-Nterm F564A | 1 ± 0.1 | 1 ± 0.1 | 61 ± 13 |
| 7TM+β_2_-Nterm L402A L403A | 6 ± 0.1 | 6 ± 0.2 | 98 ± 20 |
| Wild type | 9 ± 0.5 | 100 ± 2.6 | 100 ± 9 |
| Empty vector | 1 ± 0.1 | 1 ± 0.1 | 0 ± 2 |
| I396A | 21 ± 1 | 27 ± 0.7 | 21 ± 5 |
| I396A F564A | 1 ± 0.1 | 1 ± 0.1 | 12 ± 4 |
| I396A L402A L403A | 1 ± 0.1 | 1 ± 0.1 | 19 ± 5 |
| S397A | 55 ± 2.8 | 71 ± 2.9 | 20 ± 1 |
| S397A F564A | 1 ± 0.1 | 2 ± 0.1 | 13 ± 4 |
| S397A L402A L403A | 1 ± 0.1 | 1 ± 0.1 | 15 ± 3 |
| ΔLDLa+Linker | 13 ± 0.7 | 14 ± 0.5 | 95 ± 2 |
| ΔLDLa+Linker S397A | 77 ± 2.4 | 79 ± 1.5 | 46 ± 2 |

**Table S4. Binding and expression data for RXFP1 constructs in Fig. 4e,f.**

^†^Mean ± s.e.m., n=3 technical replicates.

| Construct | Fc-relaxin fusion binding (SE301) (%)^†^ | Cell surface expression (%)^†^ |
| --- | --- | --- |
| Wild type | 100 ± 8 | 100 ± 9 |
| E206A | 42 ± 7 | 101 ± 4 |
| Empty vector | 0 ± 4 | 0 ± 0.5 |

**Table S5. Binding and expression data for RXFP1 constructs in Fig. S5e,f.**

^†^Mean ± s.e.m., n=3 technical replicates.

| Construct | Ratio of binding to expression^†^ |
| --- | --- |
| Wild type | 0.82 ± 0.03 |
| ΔLDLa | 1.41 ± 0.04 |
| ΔLDLa+Linker | 0.16 ± 0.01 |
| GGS Linker | 0.15 ± 0.01 |
| F564A | 0.40 ± 0.01 |
| L566A | 0.63 ± 0.01 |
| Wild type | 0.62 ± 0.11 |
| S397A | 0.44 ± 0.11 |
| I396A | 0.37 ± 0.06 |
| L402A L403A | 0.37 ± 0.07 |
| 7TM+β_2_-Nterm | 0.19 ± 0.02 |

**Table S6, related to Fig. 3d. Hydrogen bonds in the hinge region disrupted by the S397A mutation.**

| H-bond (lifetime during MD, in %) | WT | S397A | S397A L402A L403A |
| --- | --- | --- | --- |
| S397 side chain – L402 backbone | 58 | 0 | 0 |
| S397 side chain – D394 side chain | 30 | 0 | 0 |
| S397 backbone – D394 backbone | 39 | 22 | 39 |

**Table S7. G_s_ signaling and expression data for RXFP1 constructs in Fig. S10.**

^†^Mean ± s.e.m., n=3 technical replicates.

See Fig.S5 and Tables S2, S3 for construct expression data.

| Construct | Basal signaling (%)^†^ | ML290 (%)^†^ |
| --- | --- | --- |
| Wild type | 13 ± 0.6 | 100 ± 1.9 |
| F564A | 1 ± 0.2 | 21 ± 2.1 |
| P565A | 1 ± 0.1 | 66 ± 8.3 |
| 7TM+β_2_-Nterm | 95 ± 2.5 | 100 ± 6.1 |
| 7TM+β_2_-Nterm F564A | 2 ± 0.2 | 2 ± 0.2 |
| 7TM+β_2_-Nterm L402A L403A | 6 ± 0.9 | 5 ± 0.4 |

| Table S8. Construct sequences | |
| --- | --- |
| Name | **Sequence** |
| SE001 | MKTIIALSYIFCLVFAHHHHHHDSWMEEVIKLCGRELVRAQIAICGMSTWSDAASSHSHSSARQLYSALANKCCHVGCTKRSLARFC |
| SE301 | MKTIIALSYIFCLVFADKTHTCPPCPAPELLGGPSVFLFPPKPKDTLMISRTPEVTCVVVDVSHEDPEVKFNWYVDGVEVHNAKTKPREEQYQSTYRVVSVLTVLHQDWLNGKEYKCKVSNKALPAPIEKTISKAKGQPREPQVYTLPPSREEMTKNQVSLTCLVKGFYPSDIAVEWESNGQPENNYKTTPPVLDSDGSFFLYSKLTVDKSRWQQGNVFSCSVMHEALHNHYTQKSLSLSPGKGGSDSWKEEVIKLCGRELVRAQIAICGKSTASDAAGANANAGARQLYSALANKCCHVGCTKRSLARFC |
| WT RXFP1 | MKTIIALSYIFCLVFADYKDDDDQDVKCSLGYFPCGNITKCLPQLLHCNGVDDCGNQADEDNCGDNNGWSLQFDKYFASYYKMTSQYPFEAETPECLVGSVPVQCLCQGLELDCDETNLRAVPSVSSNVTAMSLQWNLIRKLPPDCFKNYHDLQKLYLQNNKITSISIYAFRGLNSLTKLYLSHNRITFLKPGVFEDLHRLEWLIIEDNHLSRISPPTFYGLNSLILLVLMNNVLTRLPDKPLCQHMPRLHWLDLEGNHIHNLRNLTFISCSNLTVLVMRKNKINHLNENTFAPLQKLDELDLGSNKIENLPPLIFKDLKELSQLNLSYNPIQKIQANQFDYLVKLKSLSLEGIEISNIQQRMFRPLMNLSHIYFKKFQYCGYAPHVRSCKPNTDGISSLENLLASIIQRVFVWVVSAVTCFGNIFVICMRPYIRSENKLYAMSIISLCCADCLMGIYLFVIGGFDLKFRGEYNKHAQLWMESTHCQLVGSLAILSTEVSVLLLTFLTLEKYICIVYPFRCVRPGKCRTITVLILIWITGFIVAFIPLSNKEFFKNYYGTNGVCFPLHSEDTESIGAQIYSVAIFLGINLAAFIIIVFSYGSMFYSVHQSAITATEIRNQVKKEMILAKRFFFIVFTDALCWIPIFVVKFLSLLQVEIPGTITSWVVIFILPINSALNPILYTLTTRPFKEMIHRFWYNYRQRKSMDSKGQKTYAPSFIWVEMWPLQEMPPELMKPDLFTYPCEMSLISQSTRLNSYS |
| RXFP1 N560A | MKTIIALSYIFCLVFADYKDDDDQDVKCSLGYFPCGNITKCLPQLLHCNGVDDCGNQADEDNCGDNNGWSLQFDKYFASYYKMTSQYPFEAETPECLVGSVPVQCLCQGLELDCDETNLRAVPSVSSNVTAMSLQWNLIRKLPPDCFKNYHDLQKLYLQNNKITSISIYAFRGLNSLTKLYLSHNRITFLKPGVFEDLHRLEWLIIEDNHLSRISPPTFYGLNSLILLVLMNNVLTRLPDKPLCQHMPRLHWLDLEGNHIHNLRNLTFISCSNLTVLVMRKNKINHLNENTFAPLQKLDELDLGSNKIENLPPLIFKDLKELSQLNLSYNPIQKIQANQFDYLVKLKSLSLEGIEISNIQQRMFRPLMNLSHIYFKKFQYCGYAPHVRSCKPNTDGISSLENLLASIIQRVFVWVVSAVTCFGNIFVICMRPYIRSENKLYAMSIISLCCADCLMGIYLFVIGGFDLKFRGEYNKHAQLWMESTHCQLVGSLAILSTEVSVLLLTFLTLEKYICIVYPFRCVRPGKCRTITVLILIWITGFIVAFIPLSNKEFFKNYYGT**A**GVCFPLHSEDTESIGAQIYSVAIFLGINLAAFIIIVFSYGSMFYSVHQSAITATEIRNQVKKEMILAKRFFFIVFTDALCWIPIFVVKFLSLLQVEIPGTITSWVVIFILPINSALNPILYTLTTRPFKEMIHRFWYNYRQRKSMDSKGQKTYAPSFIWVEMWPLQEMPPELMKPDLFTYPCEMSLISQSTRLNSYS |
| RXFP1 F564A | MKTIIALSYIFCLVFADYKDDDDQDVKCSLGYFPCGNITKCLPQLLHCNGVDDCGNQADEDNCGDNNGWSLQFDKYFASYYKMTSQYPFEAETPECLVGSVPVQCLCQGLELDCDETNLRAVPSVSSNVTAMSLQWNLIRKLPPDCFKNYHDLQKLYLQNNKITSISIYAFRGLNSLTKLYLSHNRITFLKPGVFEDLHRLEWLIIEDNHLSRISPPTFYGLNSLILLVLMNNVLTRLPDKPLCQHMPRLHWLDLEGNHIHNLRNLTFISCSNLTVLVMRKNKINHLNENTFAPLQKLDELDLGSNKIENLPPLIFKDLKELSQLNLSYNPIQKIQANQFDYLVKLKSLSLEGIEISNIQQRMFRPLMNLSHIYFKKFQYCGYAPHVRSCKPNTDGISSLENLLASIIQRVFVWVVSAVTCFGNIFVICMRPYIRSENKLYAMSIISLCCADCLMGIYLFVIGGFDLKFRGEYNKHAQLWMESTHCQLVGSLAILSTEVSVLLLTFLTLEKYICIVYPFRCVRPGKCRTITVLILIWITGFIVAFIPLSNKEFFKNYYGTNGVC**A**PLHSEDTESIGAQIYSVAIFLGINLAAFIIIVFSYGSMFYSVHQSAITATEIRNQVKKEMILAKRFFFIVFTDALCWIPIFVVKFLSLLQVEIPGTITSWVVIFILPINSALNPILYTLTTRPFKEMIHRFWYNYRQRKSMDSKGQKTYAPSFIWVEMWPLQEMPPELMKPDLFTYPCEMSLISQSTRLNSYS |
| RXFP1 P565A | MKTIIALSYIFCLVFADYKDDDDQDVKCSLGYFPCGNITKCLPQLLHCNGVDDCGNQADEDNCGDNNGWSLQFDKYFASYYKMTSQYPFEAETPECLVGSVPVQCLCQGLELDCDETNLRAVPSVSSNVTAMSLQWNLIRKLPPDCFKNYHDLQKLYLQNNKITSISIYAFRGLNSLTKLYLSHNRITFLKPGVFEDLHRLEWLIIEDNHLSRISPPTFYGLNSLILLVLMNNVLTRLPDKPLCQHMPRLHWLDLEGNHIHNLRNLTFISCSNLTVLVMRKNKINHLNENTFAPLQKLDELDLGSNKIENLPPLIFKDLKELSQLNLSYNPIQKIQANQFDYLVKLKSLSLEGIEISNIQQRMFRPLMNLSHIYFKKFQYCGYAPHVRSCKPNTDGISSLENLLASIIQRVFVWVVSAVTCFGNIFVICMRPYIRSENKLYAMSIISLCCADCLMGIYLFVIGGFDLKFRGEYNKHAQLWMESTHCQLVGSLAILSTEVSVLLLTFLTLEKYICIVYPFRCVRPGKCRTITVLILIWITGFIVAFIPLSNKEFFKNYYGTNGVCF**A**LHSEDTESIGAQIYSVAIFLGINLAAFIIIVFSYGSMFYSVHQSAITATEIRNQVKKEMILAKRFFFIVFTDALCWIPIFVVKFLSLLQVEIPGTITSWVVIFILPINSALNPILYTLTTRPFKEMIHRFWYNYRQRKSMDSKGQKTYAPSFIWVEMWPLQEMPPELMKPDLFTYPCEMSLISQSTRLNSYS |
| RXFP1 L566A | MKTIIALSYIFCLVFADYKDDDDQDVKCSLGYFPCGNITKCLPQLLHCNGVDDCGNQADEDNCGDNNGWSLQFDKYFASYYKMTSQYPFEAETPECLVGSVPVQCLCQGLELDCDETNLRAVPSVSSNVTAMSLQWNLIRKLPPDCFKNYHDLQKLYLQNNKITSISIYAFRGLNSLTKLYLSHNRITFLKPGVFEDLHRLEWLIIEDNHLSRISPPTFYGLNSLILLVLMNNVLTRLPDKPLCQHMPRLHWLDLEGNHIHNLRNLTFISCSNLTVLVMRKNKINHLNENTFAPLQKLDELDLGSNKIENLPPLIFKDLKELSQLNLSYNPIQKIQANQFDYLVKLKSLSLEGIEISNIQQRMFRPLMNLSHIYFKKFQYCGYAPHVRSCKPNTDGISSLENLLASIIQRVFVWVVSAVTCFGNIFVICMRPYIRSENKLYAMSIISLCCADCLMGIYLFVIGGFDLKFRGEYNKHAQLWMESTHCQLVGSLAILSTEVSVLLLTFLTLEKYICIVYPFRCVRPGKCRTITVLILIWITGFIVAFIPLSNKEFFKNYYGTNGVCFP**A**HSEDTESIGAQIYSVAIFLGINLAAFIIIVFSYGSMFYSVHQSAITATEIRNQVKKEMILAKRFFFIVFTDALCWIPIFVVKFLSLLQVEIPGTITSWVVIFILPINSALNPILYTLTTRPFKEMIHRFWYNYRQRKSMDSKGQKTYAPSFIWVEMWPLQEMPPELMKPDLFTYPCEMSLISQSTRLNSYS |
| RXFP1 L566D | MKTIIALSYIFCLVFADYKDDDDQDVKCSLGYFPCGNITKCLPQLLHCNGVDDCGNQADEDNCGDNNGWSLQFDKYFASYYKMTSQYPFEAETPECLVGSVPVQCLCQGLELDCDETNLRAVPSVSSNVTAMSLQWNLIRKLPPDCFKNYHDLQKLYLQNNKITSISIYAFRGLNSLTKLYLSHNRITFLKPGVFEDLHRLEWLIIEDNHLSRISPPTFYGLNSLILLVLMNNVLTRLPDKPLCQHMPRLHWLDLEGNHIHNLRNLTFISCSNLTVLVMRKNKINHLNENTFAPLQKLDELDLGSNKIENLPPLIFKDLKELSQLNLSYNPIQKIQANQFDYLVKLKSLSLEGIEISNIQQRMFRPLMNLSHIYFKKFQYCGYAPHVRSCKPNTDGISSLENLLASIIQRVFVWVVSAVTCFGNIFVICMRPYIRSENKLYAMSIISLCCADCLMGIYLFVIGGFDLKFRGEYNKHAQLWMESTHCQLVGSLAILSTEVSVLLLTFLTLEKYICIVYPFRCVRPGKCRTITVLILIWITGFIVAFIPLSNKEFFKNYYGTNGVCFP**D**HSEDTESIGAQIYSVAIFLGINLAAFIIIVFSYGSMFYSVHQSAITATEIRNQVKKEMILAKRFFFIVFTDALCWIPIFVVKFLSLLQVEIPGTITSWVVIFILPINSALNPILYTLTTRPFKEMIHRFWYNYRQRKSMDSKGQKTYAPSFIWVEMWPLQEMPPELMKPDLFTYPCEMSLISQSTRLNSYS |
| RXFP1 H567A | MKTIIALSYIFCLVFADYKDDDDQDVKCSLGYFPCGNITKCLPQLLHCNGVDDCGNQADEDNCGDNNGWSLQFDKYFASYYKMTSQYPFEAETPECLVGSVPVQCLCQGLELDCDETNLRAVPSVSSNVTAMSLQWNLIRKLPPDCFKNYHDLQKLYLQNNKITSISIYAFRGLNSLTKLYLSHNRITFLKPGVFEDLHRLEWLIIEDNHLSRISPPTFYGLNSLILLVLMNNVLTRLPDKPLCQHMPRLHWLDLEGNHIHNLRNLTFISCSNLTVLVMRKNKINHLNENTFAPLQKLDELDLGSNKIENLPPLIFKDLKELSQLNLSYNPIQKIQANQFDYLVKLKSLSLEGIEISNIQQRMFRPLMNLSHIYFKKFQYCGYAPHVRSCKPNTDGISSLENLLASIIQRVFVWVVSAVTCFGNIFVICMRPYIRSENKLYAMSIISLCCADCLMGIYLFVIGGFDLKFRGEYNKHAQLWMESTHCQLVGSLAILSTEVSVLLLTFLTLEKYICIVYPFRCVRPGKCRTITVLILIWITGFIVAFIPLSNKEFFKNYYGTNGVCFPL**A**SEDTESIGAQIYSVAIFLGINLAAFIIIVFSYGSMFYSVHQSAITATEIRNQVKKEMILAKRFFFIVFTDALCWIPIFVVKFLSLLQVEIPGTITSWVVIFILPINSALNPILYTLTTRPFKEMIHRFWYNYRQRKSMDSKGQKTYAPSFIWVEMWPLQEMPPELMKPDLFTYPCEMSLISQSTRLNSYS |
| RXFP1 L402A | MKTIIALSYIFCLVFADYKDDDDQDVKCSLGYFPCGNITKCLPQLLHCNGVDDCGNQADEDNCGDNNGWSLQFDKYFASYYKMTSQYPFEAETPECLVGSVPVQCLCQGLELDCDETNLRAVPSVSSNVTAMSLQWNLIRKLPPDCFKNYHDLQKLYLQNNKITSISIYAFRGLNSLTKLYLSHNRITFLKPGVFEDLHRLEWLIIEDNHLSRISPPTFYGLNSLILLVLMNNVLTRLPDKPLCQHMPRLHWLDLEGNHIHNLRNLTFISCSNLTVLVMRKNKINHLNENTFAPLQKLDELDLGSNKIENLPPLIFKDLKELSQLNLSYNPIQKIQANQFDYLVKLKSLSLEGIEISNIQQRMFRPLMNLSHIYFKKFQYCGYAPHVRSCKPNTDGISSLEN**A**LASIIQRVFVWVVSAVTCFGNIFVICMRPYIRSENKLYAMSIISLCCADCLMGIYLFVIGGFDLKFRGEYNKHAQLWMESTHCQLVGSLAILSTEVSVLLLTFLTLEKYICIVYPFRCVRPGKCRTITVLILIWITGFIVAFIPLSNKEFFKNYYGTNGVCFPLHSEDTESIGAQIYSVAIFLGINLAAFIIIVFSYGSMFYSVHQSAITATEIRNQVKKEMILAKRFFFIVFTDALCWIPIFVVKFLSLLQVEIPGTITSWVVIFILPINSALNPILYTLTTRPFKEMIHRFWYNYRQRKSMDSKGQKTYAPSFIWVEMWPLQEMPPELMKPDLFTYPCEMSLISQSTRLNSYS |
| RXFP1 L403A | MKTIIALSYIFCLVFADYKDDDDQDVKCSLGYFPCGNITKCLPQLLHCNGVDDCGNQADEDNCGDNNGWSLQFDKYFASYYKMTSQYPFEAETPECLVGSVPVQCLCQGLELDCDETNLRAVPSVSSNVTAMSLQWNLIRKLPPDCFKNYHDLQKLYLQNNKITSISIYAFRGLNSLTKLYLSHNRITFLKPGVFEDLHRLEWLIIEDNHLSRISPPTFYGLNSLILLVLMNNVLTRLPDKPLCQHMPRLHWLDLEGNHIHNLRNLTFISCSNLTVLVMRKNKINHLNENTFAPLQKLDELDLGSNKIENLPPLIFKDLKELSQLNLSYNPIQKIQANQFDYLVKLKSLSLEGIEISNIQQRMFRPLMNLSHIYFKKFQYCGYAPHVRSCKPNTDGISSLENL**A**ASIIQRVFVWVVSAVTCFGNIFVICMRPYIRSENKLYAMSIISLCCADCLMGIYLFVIGGFDLKFRGEYNKHAQLWMESTHCQLVGSLAILSTEVSVLLLTFLTLEKYICIVYPFRCVRPGKCRTITVLILIWITGFIVAFIPLSNKEFFKNYYGTNGVCFPLHSEDTESIGAQIYSVAIFLGINLAAFIIIVFSYGSMFYSVHQSAITATEIRNQVKKEMILAKRFFFIVFTDALCWIPIFVVKFLSLLQVEIPGTITSWVVIFILPINSALNPILYTLTTRPFKEMIHRFWYNYRQRKSMDSKGQKTYAPSFIWVEMWPLQEMPPELMKPDLFTYPCEMSLISQSTRLNSYS |
| RXFP1 ΔLDLa | MKTIIALSYIFCLVFADYKDDDDGDNNGWSLQFDKYFASYYKMTSQYPFEAETPECLVGSVPVQCLCQGLELDCDETNLRAVPSVSSNVTAMSLQWNLIRKLPPDCFKNYHDLQKLYLQNNKITSISIYAFRGLNSLTKLYLSHNRITFLKPGVFEDLHRLEWLIIEDNHLSRISPPTFYGLNSLILLVLMNNVLTRLPDKPLCQHMPRLHWLDLEGNHIHNLRNLTFISCSNLTVLVMRKNKINHLNENTFAPLQKLDELDLGSNKIENLPPLIFKDLKELSQLNLSYNPIQKIQANQFDYLVKLKSLSLEGIEISNIQQRMFRPLMNLSHIYFKKFQYCGYAPHVRSCKPNTDGISSLENLLASIIQRVFVWVVSAVTCFGNIFVICMRPYIRSENKLYAMSIISLCCADCLMGIYLFVIGGFDLKFRGEYNKHAQLWMESTHCQLVGSLAILSTEVSVLLLTFLTLEKYICIVYPFRCVRPGKCRTITVLILIWITGFIVAFIPLSNKEFFKNYYGTNGVCFPLHSEDTESIGAQIYSVAIFLGINLAAFIIIVFSYGSMFYSVHQSAITATEIRNQVKKEMILAKRFFFIVFTDALCWIPIFVVKFLSLLQVEIPGTITSWVVIFILPINSALNPILYTLTTRPFKEMIHRFWYNYRQRKSMDSKGQKTYAPSFIWVEMWPLQEMPPELMKPDLFTYPCEMSLISQSTRLNSYS |
| RXFP1 ΔLDLa+Linker | MKTIIALSYIFCLVFADYKDDDDCLVGSVPVQCLCQGLELDCDETNLRAVPSVSSNVTAMSLQWNLIRKLPPDCFKNYHDLQKLYLQNNKITSISIYAFRGLNSLTKLYLSHNRITFLKPGVFEDLHRLEWLIIEDNHLSRISPPTFYGLNSLILLVLMNNVLTRLPDKPLCQHMPRLHWLDLEGNHIHNLRNLTFISCSNLTVLVMRKNKINHLNENTFAPLQKLDELDLGSNKIENLPPLIFKDLKELSQLNLSYNPIQKIQANQFDYLVKLKSLSLEGIEISNIQQRMFRPLMNLSHIYFKKFQYCGYAPHVRSCKPNTDGISSLENLLASIIQRVFVWVVSAVTCFGNIFVICMRPYIRSENKLYAMSIISLCCADCLMGIYLFVIGGFDLKFRGEYNKHAQLWMESTHCQLVGSLAILSTEVSVLLLTFLTLEKYICIVYPFRCVRPGKCRTITVLILIWITGFIVAFIPLSNKEFFKNYYGTNGVCFPLHSEDTESIGAQIYSVAIFLGINLAAFIIIVFSYGSMFYSVHQSAITATEIRNQVKKEMILAKRFFFIVFTDALCWIPIFVVKFLSLLQVEIPGTITSWVVIFILPINSALNPILYTLTTRPFKEMIHRFWYNYRQRKSMDSKGQKTYAPSFIWVEMWPLQEMPPELMKPDLFTYPCEMSLISQSTRLNSYS |
| RXFP1 GGS Linker | MKTIIALSYIFCLVFADYKDDDDQDVKCSLGYFPCGNITKCLPQLLHCNGVDDCGNQADEDNCGSGGSGGSGGSGGSGGSGGSGGSGGSGGSGGSCLVGSVPVQCLCQGLELDCDETNLRAVPSVSSNVTAMSLQWNLIRKLPPDCFKNYHDLQKLYLQNNKITSISIYAFRGLNSLTKLYLSHNRITFLKPGVFEDLHRLEWLIIEDNHLSRISPPTFYGLNSLILLVLMNNVLTRLPDKPLCQHMPRLHWLDLEGNHIHNLRNLTFISCSNLTVLVMRKNKINHLNENTFAPLQKLDELDLGSNKIENLPPLIFKDLKELSQLNLSYNPIQKIQANQFDYLVKLKSLSLEGIEISNIQQRMFRPLMNLSHIYFKKFQYCGYAPHVRSCKPNTDGISSLENLLASIIQRVFVWVVSAVTCFGNIFVICMRPYIRSENKLYAMSIISLCCADCLMGIYLFVIGGFDLKFRGEYNKHAQLWMESTHCQLVGSLAILSTEVSVLLLTFLTLEKYICIVYPFRCVRPGKCRTITVLILIWITGFIVAFIPLSNKEFFKNYYGTNGVCFPLHSEDTESIGAQIYSVAIFLGINLAAFIIIVFSYGSMFYSVHQSAITATEIRNQVKKEMILAKRFFFIVFTDALCWIPIFVVKFLSLLQVEIPGTITSWVVIFILPINSALNPILYTLTTRPFKEMIHRFWYNYRQRKSMDSKGQKTYAPSFIWVEMWPLQEMPPELMKPDLFTYPCEMSLISQSTRLNSYS |
| RXFP1 TMD | MKTIIALSYIFCLVFADYKDDDDGISSLENLLASIIQRVFVWVVSAVTCFGNIFVICMRPYIRSENKLYAMSIISLCCADCLMGIYLFVIGGFDLKFRGEYNKHAQLWMESTHCQLVGSLAILSTEVSVLLLTFLTLEKYICIVYPFRCVRPGKCRTITVLILIWITGFIVAFIPLSNKEFFKNYYGTNGVCFPLHSEDTESIGAQIYSVAIFLGINLAAFIIIVFSYGSMFYSVHQSAITATEIRNQVKKEMILAKRFFFIVFTDALCWIPIFVVKFLSLLQVEIPGTITSWVVIFILPINSALNPILYTLTTRPFKEMIHRFWYNYRQRKSMDSKGQKTYAPSFIWVEMWPLQEMPPELMKPDLFTYPCEMSLISQSTRLNSYS |
| RXFP1 TMD + β_2_-Nterm | MKTIIALSYIFCLVFADYKDDDDAMGQPGNGSAFLLAPNRSHAPDHDVTQQRGISSLENLLASIIQRVFVWVVSAVTCFGNIFVICMRPYIRSENKLYAMSIISLCCADCLMGIYLFVIGGFDLKFRGEYNKHAQLWMESTHCQLVGSLAILSTEVSVLLLTFLTLEKYICIVYPFRCVRPGKCRTITVLILIWITGFIVAFIPLSNKEFFKNYYGTNGVCFPLHSEDTESIGAQIYSVAIFLGINLAAFIIIVFSYGSMFYSVHQSAITATEIRNQVKKEMILAKRFFFIVFTDALCWIPIFVVKFLSLLQVEIPGTITSWVVIFILPINSALNPILYTLTTRPFKEMIHRFWYNYRQRKSMDSKGQKTYAPSFIWVEMWPLQEMPPELMKPDLFTYPCEMSLISQSTRLNSYS |
| RXFP1 I396A | MKTIIALSYIFCLVFADYKDDDDQDVKCSLGYFPCGNITKCLPQLLHCNGVDDCGNQADEDNCGDNNGWSLQFDKYFASYYKMTSQYPFEAETPECLVGSVPVQCLCQGLELDCDETNLRAVPSVSSNVTAMSLQWNLIRKLPPDCFKNYHDLQKLYLQNNKITSISIYAFRGLNSLTKLYLSHNRITFLKPGVFEDLHRLEWLIIEDNHLSRISPPTFYGLNSLILLVLMNNVLTRLPDKPLCQHMPRLHWLDLEGNHIHNLRNLTFISCSNLTVLVMRKNKINHLNENTFAPLQKLDELDLGSNKIENLPPLIFKDLKELSQLNLSYNPIQKIQANQFDYLVKLKSLSLEGIEISNIQQRMFRPLMNLSHIYFKKFQYCGYAPHVRSCKPNTDG**A**SSLENLLASIIQRVFVWVVSAVTCFGNIFVICMRPYIRSENKLYAMSIISLCCADCLMGIYLFVIGGFDLKFRGEYNKHAQLWMESTHCQLVGSLAILSTEVSVLLLTFLTLEKYICIVYPFRCVRPGKCRTITVLILIWITGFIVAFIPLSNKEFFKNYYGTNGVCFPLHSEDTESIGAQIYSVAIFLGINLAAFIIIVFSYGSMFYSVHQSAITATEIRNQVKKEMILAKRFFFIVFTDALCWIPIFVVKFLSLLQVEIPGTITSWVVIFILPINSALNPILYTLTTRPFKEMIHRFWYNYRQRKSMDSKGQKTYAPSFIWVEMWPLQEMPPELMKPDLFTYPCEMSLISQSTRLNSYS |
| RXFP1 S397A | MKTIIALSYIFCLVFADYKDDDDQDVKCSLGYFPCGNITKCLPQLLHCNGVDDCGNQADEDNCGDNNGWSLQFDKYFASYYKMTSQYPFEAETPECLVGSVPVQCLCQGLELDCDETNLRAVPSVSSNVTAMSLQWNLIRKLPPDCFKNYHDLQKLYLQNNKITSISIYAFRGLNSLTKLYLSHNRITFLKPGVFEDLHRLEWLIIEDNHLSRISPPTFYGLNSLILLVLMNNVLTRLPDKPLCQHMPRLHWLDLEGNHIHNLRNLTFISCSNLTVLVMRKNKINHLNENTFAPLQKLDELDLGSNKIENLPPLIFKDLKELSQLNLSYNPIQKIQANQFDYLVKLKSLSLEGIEISNIQQRMFRPLMNLSHIYFKKFQYCGYAPHVRSCKPNTDGI**A**SLENLLASIIQRVFVWVVSAVTCFGNIFVICMRPYIRSENKLYAMSIISLCCADCLMGIYLFVIGGFDLKFRGEYNKHAQLWMESTHCQLVGSLAILSTEVSVLLLTFLTLEKYICIVYPFRCVRPGKCRTITVLILIWITGFIVAFIPLSNKEFFKNYYGTNGVCFPLHSEDTESIGAQIYSVAIFLGINLAAFIIIVFSYGSMFYSVHQSAITATEIRNQVKKEMILAKRFFFIVFTDALCWIPIFVVKFLSLLQVEIPGTITSWVVIFILPINSALNPILYTLTTRPFKEMIHRFWYNYRQRKSMDSKGQKTYAPSFIWVEMWPLQEMPPELMKPDLFTYPCEMSLISQSTRLNSYS |
| RXFP1 E206A | MKTIIALSYIFCLVFADYKDDDDQDVKCSLGYFPCGNITKCLPQLLHCNGVDDCGNQADEDNCGDNNGWSLQFDKYFASYYKMTSQYPFEAETPECLVGSVPVQCLCQGLELDCDETNLRAVPSVSSNVTAMSLQWNLIRKLPPDCFKNYHDLQKLYLQNNKITSISIYAFRGLNSLTKLYLSHNRITFLKPGVFEDLHRLEWLII**A**DNHLSRISPPTFYGLNSLILLVLMNNVLTRLPDKPLCQHMPRLHWLDLEGNHIHNLRNLTFISCSNLTVLVMRKNKINHLNENTFAPLQKLDELDLGSNKIENLPPLIFKDLKELSQLNLSYNPIQKIQANQFDYLVKLKSLSLEGIEISNIQQRMFRPLMNLSHIYFKKFQYCGYAPHVRSCKPNTDGISSLENLLASIIQRVFVWVVSAVTCFGNIFVICMRPYIRSENKLYAMSIISLCCADCLMGIYLFVIGGFDLKFRGEYNKHAQLWMESTHCQLVGSLAILSTEVSVLLLTFLTLEKYICIVYPFRCVRPGKCRTITVLILIWITGFIVAFIPLSNKEFFKNYYGTNGVCFPLHSEDTESIGAQIYSVAIFLGINLAAFIIIVFSYGSMFYSVHQSAITATEIRNQVKKEMILAKRFFFIVFTDALCWIPIFVVKFLSLLQVEIPGTITSWVVIFILPINSALNPILYTLTTRPFKEMIHRFWYNYRQRKSMDSKGQKTYAPSFIWVEMWPLQEMPPELMKPDLFTYPCEMSLISQSTRLNSYS |
| RXFP1-miniG_s_399-20res | MKTIIALSYIFCLVFADYKDDDDGGSLEVLFQGPGGSQDVKCSLGYFPCGNITKCLPQLLHCNGVDDCGNQADEDNCGDNNGWSLQFDKYFASYYKMTSQYPFEAETPECLVGSVPVQCLCQGLELDCDETNLRAVPSVSSNVTAMSLQWNLIRKLPPDCFKNYHDLQKLYLQNNKITSISIYAFRGLNSLTKLYLSHNRITFLKPGVFEDLHRLEWLIIEDNHLSRISPPTFYGLNSLILLVLMNNVLTRLPDKPLCQHMPRLHWLDLEGNHIHNLRNLTFISCSNLTVLVMRKNKINHLNENTFAPLQKLDELDLGSNKIENLPPLIFKDLKELSQLNLSYNPIQKIQANQFDYLVKLKSLSLEGIEISNIQQRMFRPLMNLSHIYFKKFQYCGYAPHVRSCKPNTDGISSLENLLASIIQRVFVWVVSAVTCFGNIFVICMRPYIRSENKLYAMSIISLCCADCLMGIYLFVIGGFDLKFRGEYNKHAQLWMESTHCQLVGSLAILSTEVSVLLLTFLTLEKYICIVYPFRCVRPGKCRTITVLILIWITGFIVAFIPLSNKEFFKNYYGTNGVCFPLHSEDTESIGAQIYSVAIFLGINLAAFIIIVFSYGSMFYSVHQSAITATEIRNQVKKEMILAKRFFFIVFTDALCWIPIFVVKFLSLLQVEIPGTITSWVVIFILPINSALNPILYTLTTRPFKEMIHRFWYNYRQRKSMDSKGQKTYAPSFIWVEMWPLQEMPPELMKPDLNSKTEDQRNEEKAQREANKKIEKQLQKDKQVYRATHRLLLLGADNSGKSTIVKQMRIYHGGSGGSGGTSGIFETKFQVDKVNFHMFDVGGQRDERRKWIQCFNDVTAIIFVVDSSDYNRLQEALNLFKSIWNNRWLRTISVILFLNKQDLLAEKVLAGKSKIEDYFPEFARYTTPEDATPEPGEDPRVTRAKYFIRDEFLRISTASGDGRHYCYPHFTCAVDTENARRIFNDCRDIIQRMHLRQYELL |
| Nb35-His-PrC | MKYLLPTAAAGLLLLAAQPAMAQVQLQESGGGLVQPGGSLRLSCAASGFTFSNYKMNWVRQAPGKGLEWVSDISQSGARISYTGSVKGRFTISRDNAKNTLYLQMNSLKPEDTAVYYCARCPAPFTRDCFDVTSTTYAYRGQGTQVTVSSLEVLFQGPGHHHHHHHHGSEDQVDPRLIDGK |
| Gβ | MHHHHHHGSSGSELDQLRQEAEQLKNQIRDARKACADATLSQITNNIDPVGRIQMRTRRTLRGHLAKIYAMHWGTDSRLLVSASQDGKLIIWDSYTTNKVHAIPLRSSWVMTCAYAPSGNYVACGGLDNICSIYNLKTREGNVRVSRELAGHTGYLSCCRFLDDNQIVTSSGDTTCALWDIETGQQTTTFTGHTGDVMSLSLAPDTRLFVSGACDASAKLWDVREGMCRQTFTGHESDINAICFFPNGNAFATGSDDATCRLFDLRADQELMTYSHDNIICGITSVSFSKSGRLLLAGYDDFNCNVWDALKADRAGVLAGHDNRVSCLGVTDDGMAVATGSWDSFLKIWN |
| Gγ | MASNNTASIAQARKLVEQLKMEANIDRIKVSKAAADLMAYCEAHAKEDPLLTPVPASENPFREKKFFCAIL |

**Legend:**

Hemagglutinin signal sequence, His-tag, Human IgG1 Fc N297Q, FLAG tag, Linker residues (not RXFP1 domain)

β_2_ adrenergic receptor N-terminus, 3C protease cleavage site, miniG_s_399, pelB signal sequence, Protein C tag**, Mutations**
